# Ecological and evolutionary patterns of virus-host interactions throughout a grassland soil depth profile

**DOI:** 10.1101/2022.12.09.519740

**Authors:** George Muscatt, Ryan Cook, Andrew Millard, Gary D. Bending, Eleanor Jameson

## Abstract

**Background:** Soil microbes play pivotal roles in global carbon cycling, however the fundamental interactions between microbes and their infecting viruses remain unclear. This is exacerbated with soil depth, where the patterns of viral dispersal, ecology, and evolution are markedly underexplored. To investigate viral communities across soil depth, we leveraged a publicly available metagenomic data set sampled from grassland soil in northern California.

**Results:** 10,196 non-redundant vOTUs were recovered from soil sampled from 20 cm to 120 cm below the surface. Viral prevalence was high throughout the soil depth profile, with viruses infecting dominant soil phyla, including *Actinomycetota*. Contrary to leading hypotheses, lysogeny did not dominate in the soil viral communities. Viral diversity was investigated at both the population-level (i.e., macro diversity) and strain-level (i.e., micro diversity) to reveal diverse ecological and evolutionary patterns of virus-host interactions in surface and subsurface soil.

**Conclusions:** By investigating viral micro diversity in soil for the first time, we have uncovered patterns of antagonistic co-evolution across both surface and subsurface soils. Furthermore, we have provided evidence of soil viruses augmenting the remineralisation of soil carbon. While we continue to yield a more comprehensive understanding of soil viral ecology, our work appeals to future researchers to continue to investigate subsurface viral communities.

## Background

Soil microbes are integral members of terrestrial ecosystems, with microbial metabolism contributing to global carbon cycling [1]. As obligate parasites of microbes, viruses can control their hosts’ population size through lytic infections and influence their hosts’ metabolic potential through the expression of auxiliary metabolic genes (AMGs) [2–6]. In the oceans, where virus-host interactions have been more thoroughly studied, viral lysis is estimated to turnover ∼ 20% of microbial biomass each day [7]. The subsequent liberation of dissolved carbon and nutrients increases microbial respiration and limits trophic transfer up the food web [8, 9]. Despite an appreciation for the ecological roles of viruses in marine ecosystems, the relevant functions of viruses in terrestrial ecosystems have received less attention. To resolve this, recent methodological developments have provided the means to investigate soil viral ecology through metagenomics [10–12], and we are beginning to uncover the ecosystem-level impacts of soil viruses [13].

Integral to understanding soil viral ecology are the fundamentals of viral dispersal, prevalence, and persistence. The consequence of these factors is demonstrated by the structuring of viral communities across gradients of space [14–18], time [13, 16, 17], and root/soil compartment [13, 19]. However, most ecological studies have focussed on surface soils, rendering subsurface viral communities markedly underexplored. This is particularly alarming given the disparity in soil biogeochemistry between surface and subsurface niches. For example, more than half of terrestrial carbon stocks are sequestered in subsurface soils [20], with microbial respiration and biomass turnover dictating long-term carbon storage [21, 22]. Additionally, subsurface microbial communities are key drivers of pollutant biodegradation, thus controlling their fate and dispersal to groundwater resources [23]. Given the pressures of viral infection on the mortality and metabolism of host populations, investigations into subsurface soil ecology could inform global actions for mitigating climate change and promoting bioremediation.

Numerous physicochemical properties of soil vary throughout its vertical profile [24, 25]. These factors shape the distribution of microbial populations such that community variation with depth is comparable to the variation observed between surface soils from different biomes [26]. Thus, the structuring of microbial communities may reflect variation in microbial responses to nutrient availability between ecological niches. Given the requirement of host cellular machinery for replication and the specificity of host infection, the structuring of viral communities is likely highly dependent on that of their host community. Subsequently, there is great importance in characterising the diversity of fundamental virus-host interactions.

Amid exponentially decreasing host biomass, activity, and diversity in subsurface soil [26, 27], virus-host interactions are likely to vary considerably with depth. For example, microscopic investigations have found that virus-to-bacteria ratios decrease with soil depth [28]. Lower virus-to-bacteria ratios have been associated with an increased prevalence of lysogeny [28, 29], a latent replication strategy where the viral genome replicates passively within the host’s chromosome until induced. Lysogenic infections can have significant impacts on the ecology and evolution of their host communities (hereafter referred to as “eco-evolutionary interactions”) [30]. While temperate viruses, capable of lysogeny, have been predicted to dominate in soils [31–33], relevant metagenomic studies have failed to corroborate this [12, 13, 34]. The argument for such increased lysogeny, namely the reduced access to viable hosts [28, 29, 35], has a stronger case in subsurface soil. Therefore, more studies investigating subsurface viruses are required to determine infection strategy preferences throughout the soil depth profile.

The co-evolution of viruses and their hosts contributes to the emergence and maintenance of phenotypic diversity in both partners [36–38]. This relationship is inherently antagonistic since the adaptation of one partner disadvantages the survival of the other. However, we understand very little about in situ antagonistic co-evolution, and even less across environmental gradients such as soil depth. Given the stark differences in nutrient availability over short vertical distances [24, 25], which have been evidenced to impact co-evolution dynamics [39], we hypothesise that the eco-evolutionary interactions between viruses and their hosts vary throughout the soil depth profile. This is likely to implicate soil viruses in the major biogeochemical processes existing throughout soil, as has been demonstrated for marine ecosystems [8, 40].

In this study, we leveraged a publicly available metagenomic data set assembled from Californian grassland soil [41] to investigate viral communities from 20 cm to 115 cm below the soil surface. Grasslands cover ∼ 40% of non-glacial land area [42], store a third of global terrestrial carbon [43], and provide numerous ecosystem services from food production to erosion regulation [44]. This presents grassland ecosystems as an ideal model system for investigating the eco-evolutionary interactions between soil viruses and their microbial hosts. Two soil depth profiles were sampled, representing contrasting aboveground vegetation: under a Garry oak tree (“Garry Oak” samples) versus neighbouring grassland (“Hilly grassland” samples). To uncover patterns of viral dispersal, ecology, and evolution across soil depth, we assessed viral diversity at both the population-level (i.e., macro diversity) and strain-level (i.e., micro diversity). This study aimed to answer the following questions: (1) To what extent does soil depth shape the assembly of viral communities, and is this effect consistent between sites? (2) Does lysogeny vary throughout the soil depth profile, such that temperate viruses dominate in subsurface soil? (3) How do the eco-evolutionary interactions between viruses and their hosts vary throughout the soil depth profile?

## Methods

### Field site

Soil was sampled previously [41] at the Sagehorn study site within the Eel River Critical Zone Observatory in Northern California. The site is underlain by the Central Belt of the Franciscan Formation, a mélange of sheared argillaceous matrix containing blocks of sandstone and other lithologies [45]. The soil profile comprises a surface organic-rich horizon (∼ 30 cm) underlain by a clay-rich horizon (10 cm – 20 cm), directly above saprolite [46]. As a result of the low-porosity bedrock, the critical zone layers become entirely saturated during the winter wet season [46]. Sagehorn is primarily a grassland ecosystem, with scattered Garry oak (*Quercus garryana*) trees. The region has a Mediterranean climate, described by hot, dry summers (from May – September) and cool, wet winters. The average rainfall for the region is ∼ 1800 mm, with 1976 mm of precipitation recorded during the year that soil samples were taken [46].

### Sample collection

The collection of soil samples was previously performed at the Sagehorn study site in Northern California in June 2016, by Sharrar et al. [41]. The vertical soil depth profile was sampled at 20 cm, 40 cm, 60 cm, 80 cm, 100 cm, and 115 cm. Soil pits were dug using a jackhammer, and the walls of the pits were sampled on both sides with a sterile scoop, resulting in two samples per soil depth collected approximately 10 cm apart laterally. Soil was sampled at two sites: under a Garry oak tree (“Garry oak” samples) and from the grassland approximately 10 m away (“Hilly grassland” samples), for a total of 24 samples.

### Metagenomic data set access

The metagenomes assembled from each soil sample described above were accessed from NCBI under project accession PRJNA577476 (sample accessions SAMN13153360-SAMN13153383).

### Recovery of viral populations

Viral contigs were predicted from the pooled assembled metagenomes (PRJNA577476). Double-stranded DNA (dsDNA) and single-stranded DNA (ssDNA) viral contigs ≥ 5 kilobase pairs (kb) were predicted with DeepVirFinder v1.0 [47], VIBRANT v1.2.1 [48] and VirSorter v2.2.3 [49], using permissive viral score thresholds where relevant (≥ 0.8 for DeepVirFinder and ≥ 0.5 for VirSorter). The quality of viral contigs predicted from all three tools was assessed with CheckV v0.8.1 [50], and resulting trimmed viral sequences were annotated with DRAM v1.3 [51]. Annotated viral sequences were manually curated following the selection criteria outlined by Guo et al. [52]. Additionally, viral sequences with the most confident prediction scores from DeepVirFinder (with corresponding viral scores ≥ 0.95, *p* ≤ 0.05, and length ≥ 10 kb) and from VIBRANT (with corresponding quality scores of “high quality draft” or “complete circular”, and length ≥ 10 kb) were retained. Viral sequences were clustered into viral operational taxonomic units (vOTUs) at 95% nucleotide identity across 85% of shorter sequence [53] using anicalc.py and aniclust.py scripts [50], resulting in 10,196 vOTUs ≥ 5 kb, representing approximately species-level viral populations. Additional functional gene annotations were provided with Prokka v1.14.6 [54] using the Prokaryotic Virus Remote Homologous Groups (PHROGs) database [55].

To determine whether any recovered vOTUs represented previously isolated phage species, we clustered our vOTUs with the INfrastructure for a PHAge REference Database (INPHARED) of phage genomes (accessed February 2022) [56] using anicalc.py and aniclust.py scripts [50]. Viral sequences were considered to represent the same species when they shared 95% nucleotide identity across 85% of shorter sequence [53].

### Taxonomy of viral populations

Taxonomic assessment of vOTUs was achieved through shared protein clustering using vConTACT2 v0.9.22 [57] with the INPHARED phage genome database (accessed February 2022) [56], and otherwise default settings. The resultant genome network was visualised in R v4.0.5 [58] using ggnet2 from GGally v2.1.2 [59] and the Fruchterman-Reingold force-directed algorithm. Nodes (representing viral genomes) were connected by edges (representing shared protein homology), with significant connections forming viral clusters (VCs) representing roughly genus-level groups. Viral genomes sharing overlap with genomes from multiple VCs were considered as singletons. To further interrogate the similarity of recovered vOTUs to a database of > 600,000 environmental phage sequences, we leveraged the web-based PhageClouds tool [60], using an intergenomic distance threshold of 0.21.

The phylogeny of jumbo phage vOTU and “jumbo-related” vOTU genomes was investigated using the DNA polymerase gene. The translated DNA polymerase gene sequences were queried against the INPHARED phage genomes database [56] (accessed June 2022) to identify closely related phage genomes using the ublast command from USEARCH v10.0.240 [61] and a similarity E-value threshold < 0.001. For downstream visualisation, an outgroup of human alphaherpesvirus 1 was included in the analysis. The translated sequences of the DNA polymerase gene from the vOTUs and reference genomes were then aligned using MAFFT v7.271 [62, 63], with automated settings. Phylogenetic trees were constructed using IQ-TREE v1.6.3 [64–66], the Whelan and Goldman protein substitution model, and 1000 bootstrap replicates. Trees were subsequently visualised in R using ggtree v2.5.3 [67–69].

### Characterisation of viral populations

vOTUs were classified as temperate when they were identified by any of the three following methods. Firstly, if the viral contig was excised from a flanking host scaffold by CheckV. Secondly, vOTUs carrying at least one gene associated with lysogeny (i.e., transposase, integrase, excisionase, resolvase, and recombinase) were considered temperate. Lysogeny associated genes were identified using the Pfam domains: PF07508, PF00589, PF01609, PF03184, PF02914, PF01797, PF04986, PF00665, PF07825, PF00239, PF13009, PF16795, PF01526, PF03400, PF01610, PF03050, PF04693, PF07592, PF12762, PF13359, PF13586, PF13610, PF13612, PF13701, PF13737, PF13751, PF13808, PF13843 and PF13358, as previously described [70, 71]. Thirdly, vOTUs which formed a VC with at least one known temperate phage were also considered temperate.

Host assignment was achieved using a combination of methods. Firstly, hosts were inferred using the microbial taxonomy assigned to the scaffold from which proviral sequences were excised from. Secondly, CRISPR spacers identified from assembled scaffolds using PILER-CR v1.06 [72] were used to identify complementary protospacers among vOTU genomes using BLASTn, with default settings and allowing for ≤ 2 mismatches. Additionally, CrisprOpenDB [73] was used with default settings. Lastly, host genera were predicted *de novo* using WIsH v1.0 [74] and a null model trained against 9620 bacterial genomes, as previously described [70]. Given that some vOTUs had conflicting host predictions between methods, and that only a single host was considered per vOTU in our analyses, preferential assignment of hosts was ordered: provirus hosts > CRISPR spacer linkage to MAG > CRISPR spacer linkage to database genome > WIsH *de novo* prediction.

Putative viral-encoded AMGs were identified using DRAM-v [51]. Due to the expected increased false positive signal arising from the high non-viral sequence space in the soil metagenomes, strict curation of candidate AMGs was performed, as suggested [75]. Briefly, this included genes on viral contigs ≥ 10 kb or complete genomes, with an auxiliary score of 1 – 3, and with both the “M” flag (corresponding to metabolic function) and the “F” flag (corresponding to genes within 5000 bases of the end of the viral contig).

AMGs encoding carbohydrate-active enzymes (CAZymes) were further interrogated for the detection of conserved functional domains using the Conserved Domain Search (CD-Search) service [76, 77]. No CAZymes had the “A” flag from DRAM-v, which indicates tail-association, implicating putative CAZymes with host metabolism instead of viral attachment.

### Abundance of viral populations

vOTU abundance was estimated by mapping raw metagenome reads against vOTU genomes using BBMap [78] with a minimum alignment identity of 90%. vOTUs were only considered present in a sample if ≥ 75% of the contig length was covered ≥ 1× by reads, as recommended [53, 79]. Raw reads were normalised by vOTU genome length and library sequencing depth to generate counts per kilobase million (CPM) using the following formula: ((raw reads / genome length) / sample read depth) × 1 *e*^6^.

### Recovery of microbial populations

Microbial operational taxonomic units (OTUs) were recovered using bacterial and archaeal ribosomal protein S3 (rpS3) sequences, as previously described [41]. Briefly, rpS3 sequences were identified by searching proteins predicted from the assembled metagenomes using a custom hidden Markov model. rpS3 protein taxonomy was subsequently inferred using BLASTp to search against a database of rpS3 proteins [80] with an E-value threshold of 1 *e*^-10^. While the vast majority of OTUs were assigned to bacterial phyla, some OTUs were assigned to the archaeal phylum *Euryarchaeota* or unknown phyla (Table S2).

In addition to OTUs, previously reconstructed [41] bacterial and archaeal metagenome-assembled genome (MAG) sequences were accessed. Similarly, most of these genomes belonged to bacterial phyla (Table S3).

### Abundance of microbial populations and metagenome-assembled genomes

The abundance of OTUs and MAGs were estimated by mapping raw metagenome reads against rpS3-containing scaffolds and MAG genomes, respectively, using BBMap with a minimum alignment identity of 98%. OTUs and MAGs were only considered present in a sample if ≥ 75% of the contig length was covered. Coverage per base pair was normalised for sample sequencing depth using the following formula: (raw coverage/sample read depth) × average read depth across samples.

### Viral micro diversity

The nucleotide diversity (*π*) of viral populations and the proportion of non-synonymous to synonymous polymorphism ratio (pN/pS) of each viral gene in each sample was estimated with Metapop [81] using BAM files from read mapping (see above) and default parameters, including thresholds of > 70% genome coverage and > 10 × average read depth. The total micro diversity of each sample was calculated by averaging over bootstrapped *π* values, as previously described [82].

Genes under positive selection were identified with pN/pS ratios < 1. Genes encoding putative ABC transporters were further interrogated for the detection of conserved functional domains using CD-Search.

Consensus vOTU sequences were constructed using the most common allele from variant sites identified using inStrain v1.5.7 [83] and BAM files from read mapping. Variants were called if a site had a minimum of five viral scaffold reads. Strain-level heterogeneity was subsequently estimated by computing the pairwise ANI of these sample-specific consensus sequences. Pairwise comparisons were only considered for analysis when the genome coverage between samples was > 25%.

### Identification of anti-phage systems

Anti-phage systems were identified from MAGs using DefenseFinder [84, 85] (accessed May 2022), with default settings. Only MAGs carrying complete anti-phage systems i.e., with all genes relating to the anti-phage system detected on the scaffold, were considered.

### Data analysis and visualisation

All statistical analyses were conducted using R v4.1.3 [58]. Viral community alpha (within-sample) diversity was described with Shannon’s *H* index computed on vOTU CPM profiles with phyloseq v1.38.0 [86]. Viral community evenness was estimated with Pielou’s *J* index. Viral community beta (between-sample) diversity was described by computing a Bray-Curtis dissimilarity matrix from square root transformed vOTU CPM values, and subsequently visualised with non-metric multidimensional scaling (NMDS) ordination using vegan v2.6.2 [87]. The same method was used for microbial community beta diversity, using normalised coverage values. Permutational multivariate analysis of variance (PERMANOVA) tests and Mantel tests were also performed with vegan. Pearson’s correlation coefficients and linear regression slopes were calculated with stats v4.2.1. Differential abundance analysis was performed on raw read counts with DESeq2 v1.34.0 [88]. Genome maps in Figure S10 were visualised with gggenes v0.4.1 [89]. Fig. S5B was made with ComplexUpset v1.3.3 [90, 91]. All remaining plots were generated with ggplot2 v3.3.6 [92].

## Results

### Soil viral communities were structured with soil depth at both the population-level and strain-level

To investigate viral communities with soil depth, we leveraged a publicly available metagenomic data set sampled from grassland soil in northern California [41]. Soil samples were previously collected at six intervals between 20 cm and 115 cm below the surface, at two sites representing contrasting aboveground vegetation: under a Garry oak tree (“Garry Oak” samples), and neighbouring grassland (“Hilly grassland” samples). In total, 24 assembled metagenomes were used to recover viral populations (vOTUs) using a combination of viral prediction tools. This yielded 10,196 non-redundant vOTUs (> 5 kb), representing 9664 dsDNA viral species and 532 ssDNA viral species (Table S1), with 292 vOTUs (2.9% of total) identified as complete or high-quality viral genomes. The mean vOTU genome length was ∼ 12 kb, while 19 vOTUs had genome lengths > 200 kb (largest 415,894 bp) and represented “jumbo phages” [93], of which 18 where classified as high-quality genomes.

To estimate the similarity of recovered vOTUs with all currently available phage genomes [56], shared protein-based classification was performed using vConTACT2 [57] (Fig. S1). The resultant network contained viral clusters (VCs) representing roughly genus-level taxonomic groups (Fig. S1A). There were 4124 (42.7% of total) dsDNA vOTUs and 129 (24.2% of total) ssDNA vOTUs which formed 1310 VCs and 89 VCs, respectively (Table S1). However, only ten VCs included both our vOTUs and phage genomes that had been previously isolated, demonstrating the novel viral taxonomic diversity accessed from subsurface soil in this study. The analysis was expanded to include > 600,000 previously identified environmental viral sequences, using PhageClouds [60]. Our vOTUs had intergenomic distances < 0.21 with only 85 previously discovered viral sequences in public databases (Table S4). Of the 75 viral sequences with available metadata at the time of analysis, 74 were assembled from soil.

While only three jumbo phage vOTUs shared a VC with others (cluster 259), 63 vOTUs < 200 kb shared VCs with jumbo phage vOTUs (hereafter referred to as “jumbo-related” vOTUs). To investigate the diversity of these vOTUs further, we constructed a phylogeny of 24 DNA polymerase genes identified within the genomes of eight jumbo phage vOTUs and six jumbo-related vOTUs (Fig. S2). This revealed that the vOTUs belonged to six distinct phylogenetic groups, which we denoted A-F. Further investigation of the groups with the closest known relatives (groups A, B, and F) identified that the most similar DNA polymerase genes were carried by genomes < 200 kb, therefore representing non-jumbo phages (Fig. S3).

To characterise the role of soil depth in shaping the assembly of viral communities, we assessed population-level viral diversity with soil depth (Fig. 1). This revealed that viral richness (measured through the detection of vOTUs), viral evenness (measured with Pielou’s *J* index), and viral diversity (measured with Shannon’s *H* index) significantly increased with soil depth in Garry Oak (Fig. 1A). In contrast, viral richness decreased with soil depth in Hilly grassland, where no linear relationship was observed with viral evenness and diversity (Fig. 1A). Next, we tested whether soil depth was an ecological driver of viral community composition through NMDS ordination and a PERMANOVA test. Bray-Curtis dissimilarities were structured with soil depth (*R*^2^= 0.156, *F* = 7.37, *p* = 0.002) (Fig. 1B), such that significant distance-decay relationships were observed at both sites (Fig. 1C). Additionally, viral communities were distinct between sites, with aboveground vegetation explaining more than twice the variation as soil depth (*R*^2^= 0.399, *F* = 18.8, *p* = 0.001) (Fig. 1B).

**Fig. 1:**
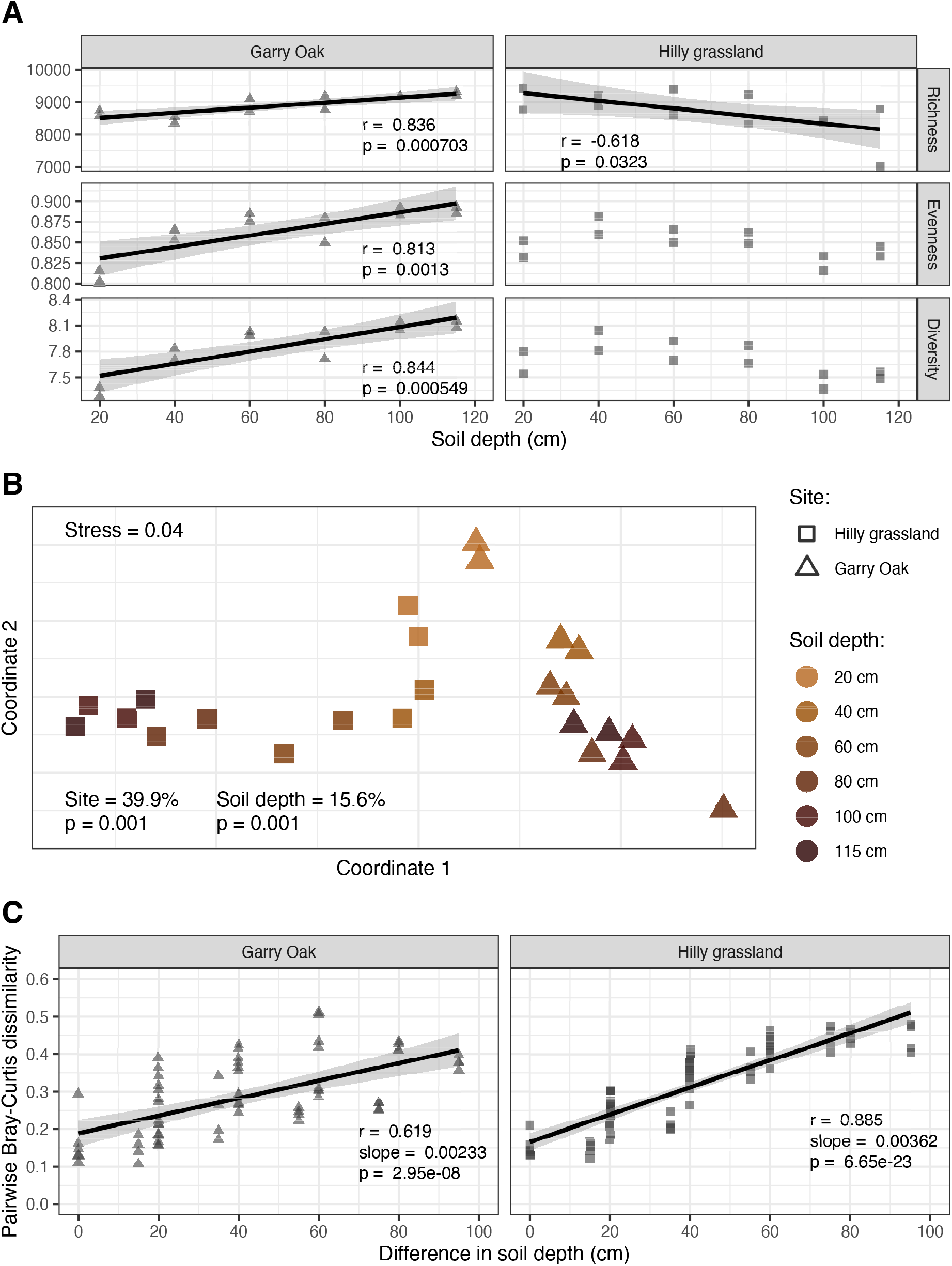
Population-level assembly of soil viral communities throughout soil depth. **A** Alpha diversity of viral communities. Richness (number of vOTUs detected), evenness (Pielou’s *J* index), and alpha diversity (Shannon’s *H* index) for each viral community throughout the soil depth profiles. Trend lines represent linear regression estimates, with shaded cloud representing 95% confidence interval. *r* corresponds to Pearson’s correlation coefficient and *p* corresponds to the associated p-value. **B** Beta diversity of viral communities. Non-metric multidimensional scaling (NMDS) ordination plots, representing the Bray-Curtis dissimilarities between viral community compositions. Shapes indicate site: Hilly grassland (squares) and Garry Oak (triangles). Shapes are coloured based on soil depth. Stress value associated with two-dimensional ordination is reported. Percentage contribution to variance by site and soil depth, as calculated with a permutational multivariate analysis of variance (PERMANOVA) test, and associated p-value are also reported. **C** Distance-decay relationship in viral community structure. Trend lines represent linear regression estimates, with shaded cloud representing 95% confidence interval. *r* corresponds to Pearson’s correlation coefficient, slope corresponds to linear regression slope, and *p* corresponds to the associated p-value.

To further contrast soil depth patterns between sites, we assessed viral prevalence to identify populations enriched in either surface or subsurface soil. This determined that viral prevalence was high throughout the soil depth profiles, such that 66.0% and 72.1% of vOTUs were shared across all samples within Garry Oak and Hilly grassland, respectively (Fig. S4). Nonetheless, differential abundance analysis identified that > 29% of vOTUs were enriched in either surface soil (20 cm) or subsurface soil (40 cm − 115 cm) (Table S1). In comparing the relative abundance of enriched viral populations between the two sites, we found that the vOTUs highly abundant in subsurface soil in one site were consistently lowly abundant throughout the soil depth profile in the other site (Fig. S5A). Subsequently, only 11.7% of depth-enriched viral populations were enriched in both sites, with 64.9% of these populations surface-enriched (Fig. S5B). In fact, subsurface-enriched viral populations in each site were genetically different, as the shared populations represented only 18.5% of subsurface-enriched VCs in Garry Oak (Fig. S1B) and 13.5% in Hilly grassland (Fig. S1C). Together, these results outline the increased distinction of subsurface soil viral communities between sites.

Lastly, we investigated the effect of soil depth in driving patterns of strain-level viral diversity (Fig. 2). To achieve this, consensus sequences were reconstructed for each vOTU in each sample, based on the most common alleles detected across variant sites. Subsequent distance-decay relationships were observed across strains of 69 vOTUs, for which the pairwise ANI between consensus sequences decreased towards 0.95 (the threshold for vOTU clustering) with soil depth (Fig. 2A). To summarise the micro diversity across viral populations of each sample, average nucleotide diversity (*π*) was assessed. This summarises the frequency of nucleotide differences between the individual strains of a population. *π* was greatest in surface soil and displayed a non-linear relationship with soil depth (Fig. 2B). As a result, no significant relationship was observed between population-level diversity (i.e., macro diversity) and strain-level diversity (i.e., micro diversity) in either site (Fig. 2C).

**Fig. 2:**
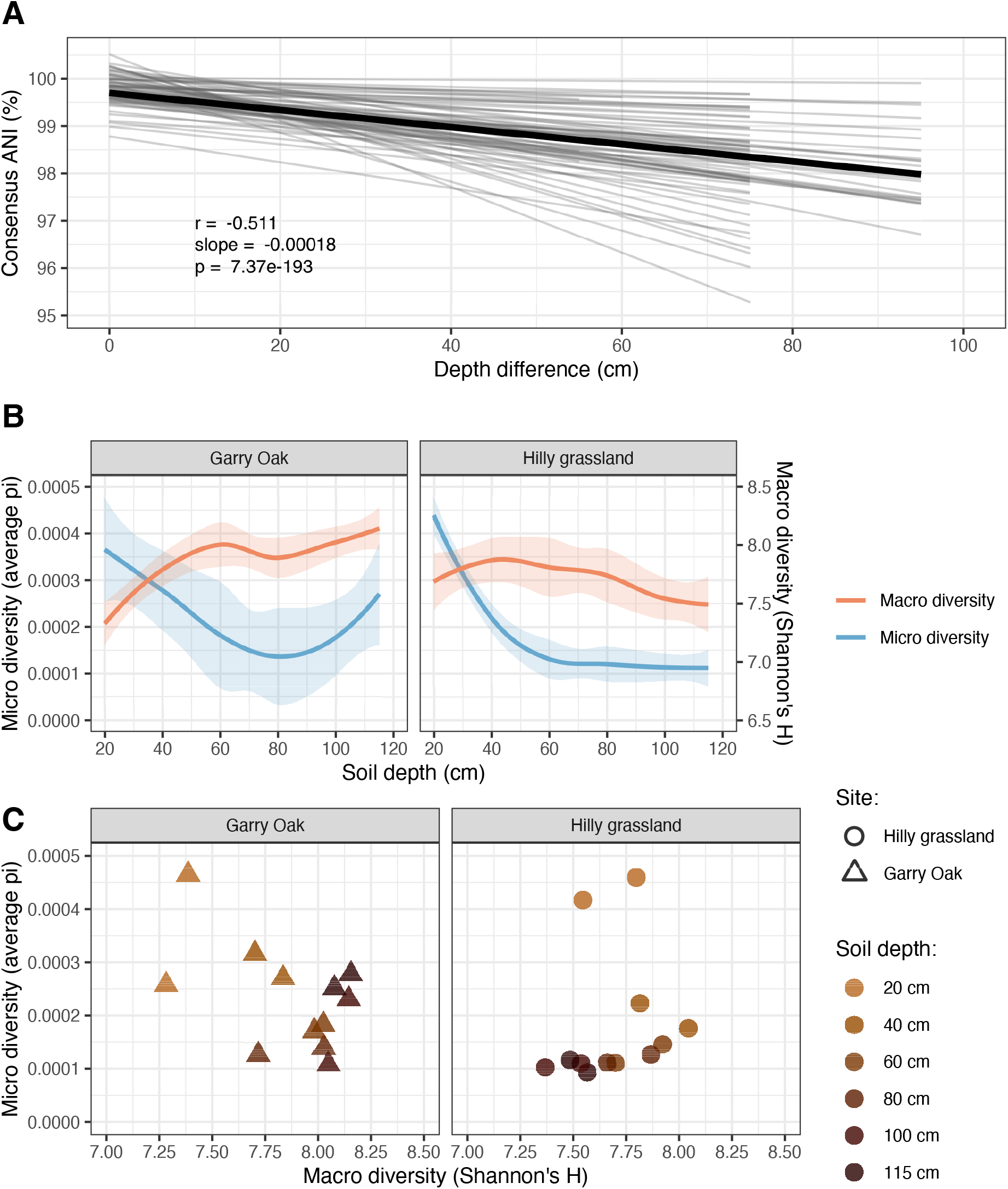
Strain-level assembly of soil viral communities throughout soil depth. **A** Distance decay relationship in consensus ANI. Lighter grey lines represent distance-decay relationships in consensus ANI for 69 vOTUs with individual significant relationships. Thicker black line represents the mean distance decay relationship across all 69 vOTUs. Trend lines represent linear regression estimates, with shaded cloud representing 95% confidence interval. *r* corresponds to Pearson’s correlation coefficient, slope corresponds to linear regression slope, and *p* corresponds to the associated p-value. **B** Viral macro diversity and micro diversity throughout the soil depth profiles. Trend lines represent loess smooth regression estimates, with shaded cloud representing 95% confidence interval. Colour indicates level of diversity: macro diversity (red), micro diversity (blue). **C** Correlation of macro diversity with micro diversity. Shapes indicate site: Hilly grassland (squares) and Garry Oak (triangles). Shapes are coloured based on soil depth.

### Virus-host interactions were diverse with soil depth

To understand the ecological role of soil viruses with the soil depth gradient, we characterised the interactions between viruses and their microbial host communities (Fig. 3). Strong links were revealed between viruses and microbes by observing significant correlations between their community structures (Fig. S6) and diversities (Fig. S7). To provide further evidence of virus-host linkages, we identified the putative host taxa of vOTUs using a combination of proviral scaffold assessment, CRISPR spacer matches, and *de novo* prediction using a probabilistic model [74]. *Actinomycetota* and *Pseudomonadota* were the most common host phyla (Table S1). Moreover, viruses infecting *Actinomycetota* were dominant members of viral communities throughout the soil depth profile of both sites (Fig. 3A). While the patterns of microbial phyla described using OTUs and MAGs were different, they both demonstrated that *Actinomycetota* abundance increased with depth in Hilly grassland (Fig. 3A). Subsequently, *Actinomycetota* and *Pseudomonadota* hosts were significantly correlated with their infecting viruses in Hilly grassland (Fig. S8).

**Fig. 3:**
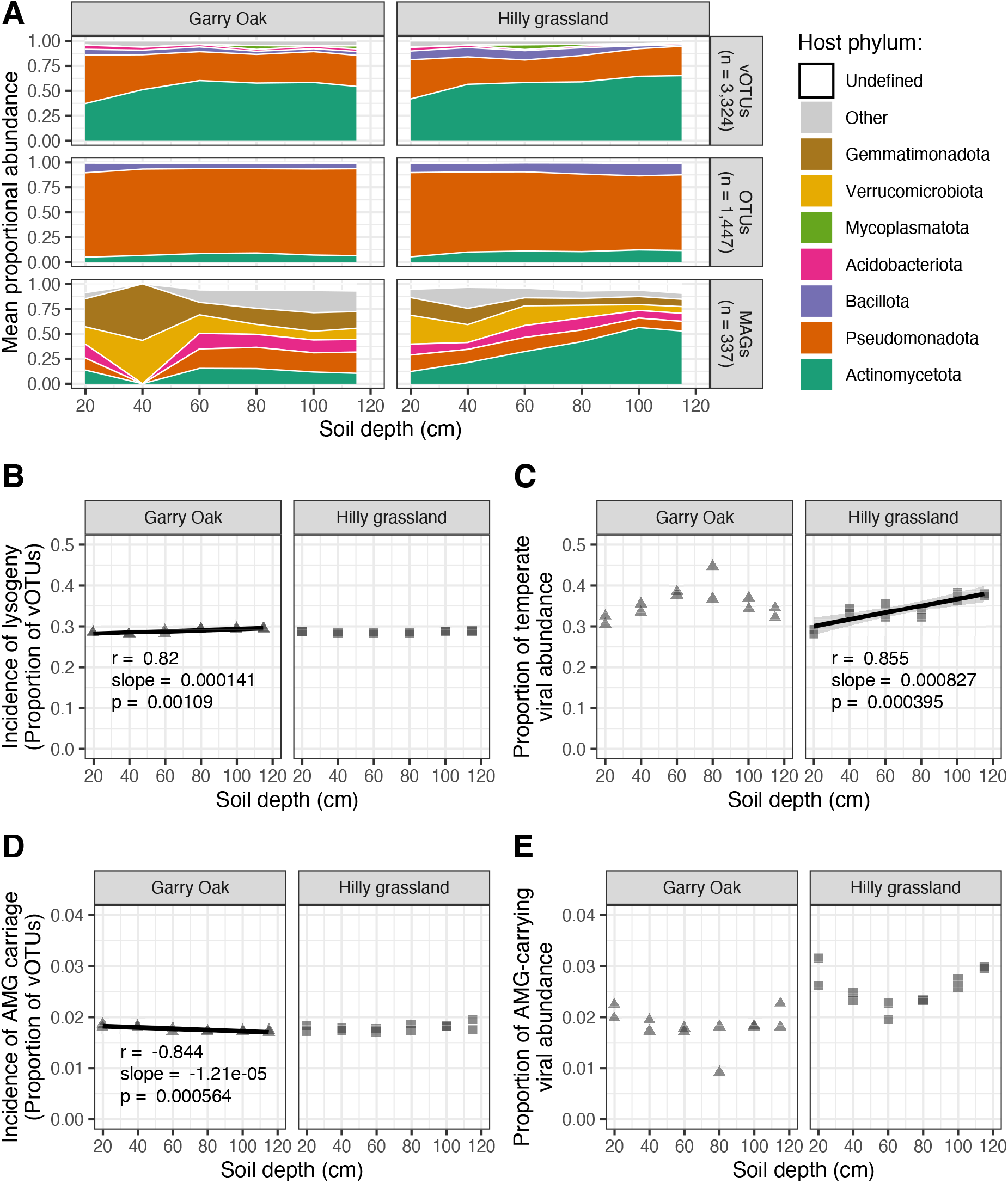
Virus-host interactions throughout soil depth. **A** Virus-host linkages. Mean proportional abundance by host phyla is plotted across soil depth for: vOTUs with predicted host phyla (n = 3324), microbial OTUs (n = 1447), and microbial MAGs (n = 337). Fill colour indicates host phylum. **B** Incidence of lysogeny. Proportion of vOTUs detected representing temperate viruses plotted across soil depth. **C** Temperate viral abundance. Proportional abundance of vOTUs detected representing temperate viruses plotted across soil depth. **D** Incidence of AMG carriage. Proportion of vOTUs carrying AMGs plotted across soil depth. **E** AMG-carrying viral abundance. Proportional abundance of vOTUs carrying AMGs plotted across soil depth. For **B**, **C**, and **D**, trend lines represent linear regression estimates, with shaded cloud representing 95% confidence interval. *r* corresponds to Pearson’s correlation coefficient, slope corresponds to linear regression slope, and *p* corresponds to the associated p-value.

Given that viral replication strategies inform virus-host interactions following infection, we investigated the prevalence of lysogeny with soil depth. In total, 2911 (28.6% of total) temperate viruses were detected. The incidence of lysogeny, as measured by the proportion of detected vOTUs which were identified as temperate, was stable across soil depth (Fig. 3B). In contrast, the relative abundance of temperate viruses varied, such that a positive relationship with soil depth was observed in Hilly grassland (Fig. 3C).

In addition to host cell lysis, another fundamental ecological role of viruses is the alteration of host metabolism through the expression of AMGs during infection. We identified 220 putative AMGs carried by 181 vOTUs (1.77% of total; Table S5), whose functional annotations included hits to ribosomal proteins (nine genes) and carbohydrate-active enzymes (CAZymes; 43 genes). Six jumbo phage vOTUs carried a single AMG each, while the average length of vOTUs carrying multiple AMGs was 29,600 bp. vOTUs carrying AMGs were consistently detected throughout the soil depth profiles, with a small yet statistically significant decrease in incidence with depth in Garry Oak (Fig. 3D). No significant depth relationships were observed for the relative abundance of AMG-carrying vOTUs (Fig. 3E).

Further inspection of candidate CAZymes with CD-Search revealed that 36/43 (83.7%) gene products contained conserved protein domains associated with carbohydrate metabolism (Table 1). This included 12 genes with glycoside hydrolase domains, putatively involved in the metabolism of four different carbon sources: glycans (five genes), amylose (two genes), cellulose (two genes), and mannose (one gene). vOTUs carrying CAZymes were dispersed across 21 VCs and 17 singletons in the shared protein network (Fig. S1D). Three quarters of vOTUs carrying CAZymes were lytic and 17/40 (42.5%) had predicted hosts, spanning *Actinomycetota* (20%), *Pseudomonadota* (12.5%), *Acidobacteriota* (5%), *Bacillota* (2.5%), and *Nitrospirota* (2.5%). The vOTUs were detected throughout the two soil depth profiles, at consistently low abundance (Fig. S9).

### Virus-host antagonistic co-evolution was dynamic throughout the soil depth profile

Virus-host interactions can also have implications on the eco-evolutionary dynamics of both viruses and microbes. Thus, to investigate virus-host antagonistic co-evolution throughout the soil depth profile, we detected bacterial anti-phage defence systems and estimated the subsequent selection pressure applied to soil viruses (Fig. 4). More than 75% of microbial community abundance was represented by MAGs carrying at least one complete anti-phage system, with systems involving restriction-modification (RM) being the most common (Fig. 4A). Further investigation into the anti-phage system repertoire of MAG communities revealed a significant increase in system diversity with soil depth in both sites (Fig. 4B).

**Fig. 4:**
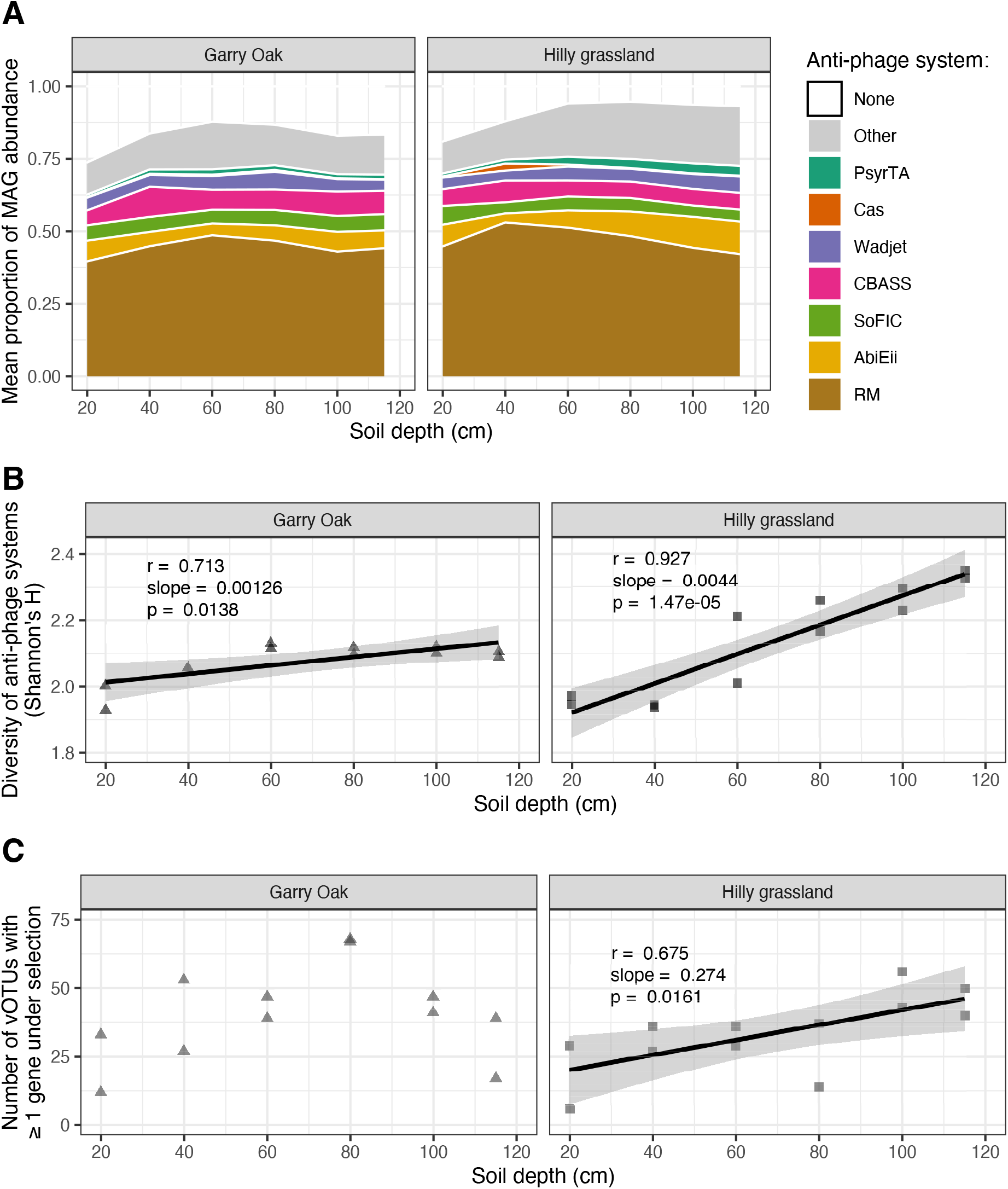
Virus-host antagonistic co-evolution throughout soil depth. **A** Anti-phage system detection. Proportional abundance of microbial MAGs carrying complete anti-phage systems. Fill colour indicates anti-phage system. **B** Diversity of the anti-phage system repertoire. Shannon’s *H* index, calculated on MAGs carrying complete anti-phage systems, plotted across soil depth. **C** Viruses under positive selection. Number of vOTU genomes with at least one gene under positive selection (indicated by a pN/pS ratio > 1) plotted across soil depth. For **B** and **C**, trend lines represent linear regression estimates, with shaded cloud representing 95% confidence interval. *r* corresponds to Pearson’s correlation coefficient, slope corresponds to linear regression slope, and *p* corresponds to the associated p-value.

To assess the resulting evolutionary pressures on viral populations, we identified viral genes under positive selection using a proportion of non-synonymous to synonymous polymorphism ratio (pN/pS) > 1. This yielded 532 vOTUs carrying 880 genes under positive selection in at least one sample, with nearly half of these genes lacking functional annotations (Table S6). Nonetheless, we were able to identify functions for 30 tail fibre proteins involved in host cell recognition [94, 95], four tape measure proteins involved in virion assembly [96] and genome insertion [97], six ribosomal proteins, and 11 ABC transporters (Table S6). Manual inspection of putative ABC transporter genes with CD-Search indicated the presence of conserved secondary structures for ten of the genes, with five genes containing drug efflux transporter domains (*ccmA*, *drrA*, *MacAB*, *MacB*, *SunT*). Moreover, five vOTUs carrying ABC transporter genes represented high-quality temperate viral genomes, with hits to viral protein families (PHROGs) both upstream and downstream of putative transporter genes (Fig. S10). While only one ABC transporter gene was positively selected in surface soil (20 cm), the remaining ten genes were positively selected in subsurface soil (40 cm – 115 cm). Overall, the number of vOTUs carrying at least one gene under positive selection increased with soil depth in Hilly grassland, while a non-linear relationship was observed with soil depth in Garry Oak (Fig. 4C).

## Discussion

### High viral dispersal maintains virus-host co-existence throughout the soil depth profile

Microbial dispersal underpins soil ecology and evolution [98], however we lack understanding of the distribution patterns of soil viruses. In this study, we observed high viral prevalence throughout two soil depth profiles, with more than two thirds of viral populations detected in every soil sample (Fig. S4). This cosmopolitan distribution contrasted with recent investigations of soil viral dispersal, in which fewer viruses were shared between samples across horizontal [14, 16, 17, 99, 100] and vertical space [18, 99, 101]. Despite high viral prevalence, we discovered that soil depth shaped the composition of viral communities (Fig. 1B), such that viral community diversity displayed a distance-decay relationship (Fig. 1C).

The structuring of viral communities with soil depth is undoubtedly driven by the physical structure of the soil matrix, which renders virion dispersal a mostly stochastic process [98]. The rate-limiting factors underlying the transport of viruses through soil are likely different to those of their hosts [32, 102]. Notably, soil viruses are expected to be passively distributed with water more easily [103]. Therefore, wetter soils may facilitate the enhanced mobility of viruses compared to their hosts, resulting in the increased accessibility and infection of susceptible host cells. Simultaneously, the abundance of viruses are also correlated with soil moisture content [14, 16, 17, 99], demonstrating how environmental factors may affect virus-host interactions.

At the Sagehorn site where soil samples were taken, significant winter precipitation raises the water table close to the soil surface [104]. The resulting annual saturation of soil may facilitate the immigration of infective viruses and susceptible hosts throughout the soil depth profile. This would have consequences on both viral and bacterial persistence due to evolutionary “source-sink dynamics”, where co-existence is maintained by the heterogeneous distribution of viruses and hosts [105, 106]. This has been demonstrated in biofilm simulations, whereby the mobility of viruses is a key determinant of phage-bacteria co-existence [107]. Therefore, we propose that the high viral dispersal is likely to have implications on the eco-evolutionary interactions occurring across the soil niches examined in this study.

### Tree association impacts viral community composition in both surface and subsurface soil

Intriguingly, the variation in viral communities between sites was greater than the variation associated with soil depth, such that communities in subsurface soils were more distinct than those at the surface (Fig. 1B). A considerable distinction between the two sites was the presence of Garry Oak trees. At the Garry Oak site, the tree canopy could have provided the soil surface with protection from the sun, potentially maintaining greater soil moisture content as compared to the unshaded soil in Hilly grassland. While changes to moisture content would be likely to affect viral dispersal and the structuring of soil viral communities, no soil property measurements were available to confirm this hypothesis.

Another consequence of Garry Oak trees is the annual shedding of leaves during winter [46]. Decaying leaf litter has been shown to shape the composition of RNA viral communities in both the rhizosphere and bulk soil [108]. While quicker degradation rates mean that the spatial structuring of RNA viruses may be greater than for DNA viruses, the legacy effects of leaf litter may have driven differences between surface soils. However, the degradation of shed leaves would be expected to have less impact on subsurface communities. Instead, we hypothesise that the presence of tree roots and the associated fungal hyphae impact viral communities in Garry Oak samples, leading to the discrepancies in the depth patterns between the two sites. Indeed, fine roots and hyphae have been reported to a depth of at least 2 m at the same study site [46]. The consequence of growing crop roots on the structures of both DNA and RNA soil viral communities has been demonstrated previously [13].

### The prevalence of lysogeny was consistent throughout the soil depth profile

Lysogenic viral infections can have significant eco-evolutionary impacts on host communities [30], most notably through superinfection exclusion, which confers resistance against further viral infection [109–111]. Typically, lysogeny is expected to dominate in soil ecosystems because of low host biomass and viability [28, 29, 35]. Under low bacterial densities (e.g., < 10^5^ cells per gram), host starvation represses viral lytic genes through ATP-dependant signalling cascades [112, 113], promoting lysogeny switching [114]. Subsequently, lower bacterial abundances have been associated with increased lysogeny in the deep ocean [115–117]. Recent work has observed an increased prevalence of lysogeny in subsurface soils, as detected through inducible lysogens [28], however we observed very little change in the incidence of temperate viruses across soil depth (Fig. 3B). And while the relative abundance of temperate phages did increase with soil depth in Hilly grassland, this was not consistent in Garry Oak (Fig. 3C). Therefore, there could be additional factors which govern lysogeny switching in soils beyond host density. This could include non-linear relationships with host metabolism [114], viral-viral interactions [118, 119], and anti-phage defence systems [85]. To this point, the diversity of anti-phage defence systems was enriched among subsurface communities in Hilly grassland (Fig. 4B), coinciding with the increased abundance of temperate viruses. The increased encountering of lysogenic infection mechanisms may have been responsible for the greater range of defence systems maintained among the host community [85]. It must also be noted that viruses without lysogenic genes can establish passive co-existence typified by temperate lifestyles, as demonstrated with ΦcrAss001 in continuous culture with its host Bacteroides intestinalis [120]. Therefore, non-lysogenic phages may be able to replicate without eradicating their host population, in contrast to the traditional view of predator-prey cycles induced by lytic phages.

### Jumbo phages recovered from soil were polyphyletic

We recovered 19 vOTUs representing jumbo phages [93] with genome lengths > 200 kb (largest 415,894 bp), without implementing a viral contig binning approach. An additional 63 vOTUs formed roughly genus-level VCs with jumbo phages, and together they represented six distinct clades based on DNA polymerase gene phylogeny (Fig. S2). This is consistent with previous findings that jumbo phages are polyphyletic, implying that phage genome gigantism has evolved numerous times instead of originating from a single common ancestor [121, 122]. Furthermore, the phylogeny revealed that the closest known relatives to jumbo phage vOTUs had much shorter genomes (Fig. S3). It has been postulated that jumbo phages may have evolved from recombination events between multiple smaller phage genomes [121]. Another potential hypothesis for the origin of phage genome gigantism is that the genomes could have expanded upon the acquisition of additional phage or host genes. The ratchet model describes how mutations that increase the capsid size facilitate the acquisition of new viral genes, which are then stable against loss of function mutations [123].

Previously identified clades of jumbo phages have been discerned by their diverse infection and replication strategies, biogeography, and host taxa [121, 122]. We have uncovered the ubiquity of jumbo phages across soil depth, suggesting that large genome sizes are evolutionarily stable across both surface and subsurface soil niches. Furthermore, jumbo phages were consistently in the top 20% of the most abundant viruses in each community (Fig. S11), contrasting with previous findings that giant viruses (> 300kb) are lowly abundant in forest soil [124].

### Soil viruses augment microbial metabolism in subsurface soils

Viruses can carry and express AMGs during infection to modulate the host’s metabolism and fitness, and promote their co-existence [2–6]. Moreover, viral-encoded AMGs have the potential to affect soil biogeochemistry, with viruses previously implicated in soil carbon processing [13, 18, 19, 34, 99, 125]. In this study, we detected viruses throughout the soil depth profile carrying CAZymes associated with both carbohydrate anabolism and catabolism (Table S5). The rank abundance of CAZyme-carrying viruses was highly variable, but their presence was ubiquitous across all soil depths (Fig. S12). Therefore, soil viruses may stimulate the degradation of a variety of carbon sources, including plant cell walls, thus contributing to the remineralisation of soil carbon in surface and subsurface soil. While our discovery of viral CAZymes adds to the repertoire of potential viral mechanisms contributing to soil carbon cycling, evidence of their function during the infection cycle has not been confirmed here.

Previously, the abundance of viral-encoded AMGs was found to increase with soil depth [101]. However, we observed that the abundance of viruses carrying AMGs was consistently low throughout both soil depth profiles (Fig. 3D-E). The most common host phyla of viruses carrying AMGs was *Actinomycetota*, for which both the host (Fig. 3A) and infecting viruses (Fig. S13) were more abundant in subsurface soil. *Actinomycetota* (formerly *Actinobacteria*) are dominant soil microbes [126] and contribute to soil carbon cycling by producing extracellular hydrolytic enzymes which depolymerise plant-derived lignin [127]. Furthermore, *Actinomycetota* are resilient to soil drying, such that their relative abundance increases during drought and declines in the days following re-wetting [128–130]. The abundance and activity blooms in response to seasonal wetting and drying are likely to affect soil nutrient and carbon cycling [130].

### Viral macro diversity and micro diversity were associated in surface soil only

The evolution of viral communities can be monitored through micro diversity. In this study, we have revealed patterns of viral micro diversity across a soil environmental gradient for the first time. Viral strain-level heterogeneity displayed a distance-decay relationship (Fig. 2A) and the average micro diversity (*π*) of viral communities varied across space (Fig. 2B).

Micro diversity is accrued through *de novo* mutations, and can drive phenotypic variation to specialise organisms to their environment [83]. More specifically for viruses, micro diversity reflects evolutionary responses to host infection dynamics, and is directly related to viral infection rates. Greater viral micro diversity, as measured by larger *π* values, can arise in multiple ways [81]. Firstly, the active infection of hosts can result in population expansion and thus more frequent mutations. This can be exacerbated through genetic recombination between viral populations co-infecting the same host. Such horizontal gene transfer events are made more likely by the presence of microbial “hotspots” occurring throughout the spatially structured soil matrix [131]. Secondly, viral populations could maintain greater micro diversity in their populations as an evolutionary mechanism. Genetic diversity increases the fitness of a viral population by allowing them to “bet-hedge” if their environment or host changes, conferring local adaptation [132].

The ecological forces driving strain-level variation were distinct from those driving population-level variation, as demonstrated by their non-significant association (Fig. 2C). This was surprising given that genetic heterogeneity between strains can result in speciation events [132, 133], thus relating the two levels of diversity. Throughout ocean depth profiles, a similar absent relationship was explained by interactions with bacterial macro diversity [82]. However, no such relationship was observed in these soil samples (Fig. S14). We speculate that unmeasured physicochemical properties, distinct between soil horizons, may have driven the non-linear diversity dynamics we observed throughout the soil depth profile.

Interestingly, when the analysis of viral diversity patterns was focussed on the top 60 cm of soil, viral macro diversity was negatively associated with viral micro diversity (Fig. 2B). This could have resulted from decreasing host cell density from surface to subsurface soil [26], which favours inter-specific viral competition (i.e., reflected in macro diversity) over intra-specific viral competition (i.e., reflected in micro diversity). Hence, strain-level heterogeneity is less favoured when fewer hosts are available, during which species-level competition drives evolution. This would be expected to impact virus-host interactions by reducing the resilience of the subsurface soil niche.

### Antagonistic co-evolution was distinct among surface and subsurface communities

Host defence responses to viral infection are expected to drive positive selection among soil viruses through antagonistic co-evolution. To this aim, we identified 880 viral genes under positive selection (Table S6), for which non-synonymous polymorphisms were more likely to be retained than rejected. This included 30 tail fibre genes, which have previously been shown to be positively selected among gut phages as evidence of their adaptive evolution [134, 135]. Phage tail fibre proteins are involved in host tropism [94, 95], thus the carriage of genetically diverse tail fibre genes may expand a population’s host range. Given the positive selection of tail fibre gene mutants throughout the soil depth profile, the evolutionary benefit of expanding host range was universal among viruses occupying both surface and subsurface soil niches.

We also identified 11 ABC transporter genes under positive selection, predominantly in subsurface soil (40 cm – 115 cm) (Table S6). Five vOTUs carrying ABC transporter genes represented high-quality temperate viral genomes (Fig. S10), with two of these genes sharing conserved protein domains with ABC drug efflux transporters. By expressing these genes during infection, temperate soil viruses may confer antibiotic resistance to their hosts, thus maintaining their mutual co-existence. Furthermore, the evidence of adaptive evolution among these genes indicates that there is a selection pressure on these viruses to augment their host’s interbacterial competition. While this may be the first evidence of soil viruses carrying ABC transporters, the expression of phosphate-binding *pstS* genes by cyanophages has implicated marine viruses in enhancing phosphate uptake in cyanobacterial hosts [136]. Many other viral genes under positive selection had no functional annotation, suggesting that we may be missing alternative selection pressures on soil viruses. For example, missing annotations may include uncharacterised anti-defence proteins, expressed by viruses to target host defence systems and maintain infective capabilities [137].

To characterise the range of host defence responses to viral infection, we identified anti-phage defence systems within microbial MAGs. The relative abundance of MAGs adopting at least one system was high throughout the soil depth profile (Fig. 4A), and the increasing diversity of anti-phage systems (Fig. 4B) suggested that the antagonistic co-evolution landscape differed between surface and subsurface niches. Multiple anti-phage defence systems can be carried within defence islands [138], a genetic toolbox of diverse mechanisms to resist viral infection, presumably accrued through horizontal gene transfer events [137]. The genetic diversity of infecting viruses can direct the evolution of host defence strategies, such that low viral diversity may favour CRISPR-based immunity, while higher viral diversity promotes surface modification mechanisms [139]. Thus, the microheterogeneity driven by the soil matrix would make these virus-host interactions difficult to predict.

## Conclusions

Most soil viral ecology efforts have focussed on the top 20 cm of soil, hindering our understanding of subsurface viruses. Given the exponential decay in microbial biomass with soil depth, one might expect relatively minimal ecological impacts of subsurface viral communities. To the contrary, we have uncovered evidence of soil viruses contributing to terrestrial ecology in both surface and subsurface soil niches. The prevalence of lysogeny was consistent throughout the soil depth profile, indicating that additional factors beyond host cell density may govern lysogeny switching in soils. By investigating patterns of viral micro diversity across a soil environmental gradient for the first time, we revealed that the local adaptation of viruses was greatest in surface soil. Furthermore, an increasing diversity of anti-phage defence systems with depth suggests that the antagonistic co-evolution landscape is distinct in subsurface soil. In the future, we predict that comparative activity studies, contrasting surface and subsurface niches, will be essential to characterise viral functions associated with soil depth.

## Abbreviations

AMG: Auxiliary Metabolic Gene
CAZyme: Carbohydrate-Active enZymes
CD-Search: Conserved Domain Search
CPM: Counts Per kilobase Million
dsDNA: double-stranded DNA
kb: kilobases
MAG: Metagenome-Assembled Genome
NMDS: Non-metric Multi-Dimensional Scaling
OTU: Operational Taxonomic Unit
PERMANOVA: PERmutational Multivariate ANalysis Of Variances
pN/pS: proportion of Non-synonymous to Synonymous polymorphism ratio
RM: Restriction Modification
rpS3: ribosomal protein S3
ssDNA: single-stranded DNA
VC: Viral Cluster
vOTU: Viral Operational Taxonomic Unit.

## Declarations

### Ethics approval and consent to participate

Not applicable.

### Consent for publication

Not applicable.

### Availability of data and materials

The metagenomic data set can be accessed from NCBI under project accession PRJNA577476 (sample accessions SAMN13153360-SAMN13153383). DNA vOTU genome sequences were deposited to the European Nucleotide Archive (ENA) under project accession PRJEB57765 (sample accession SAMEA112154074). FASTA nucleotide files containing vOTU genomes, FASTA amino acid files containing vOTU genes, vOTU gene annotations, vConTACT2 network input and output files, rpS3 protein sequences, and assembled MAG sequences are available from figshare (https://figshare.com/XXX). The custom R script used to generate figures and tables, along with required input files, are available from GitHub (https://github.com/GeorgeMuscatt/GrasslandDepthVirome).

### Authors’ contributions

G.M., A.M., G.D.B. and E.J. conceived and designed the analyses. G.M. accessed the data set, carried out bioinformatic analyses, generated R scripts, interpreted data, prepared figures, and produced the first draft of the manuscript. R.C. aided with bioinformatic analyses. A.M., G.D.B., and E.J. provided edits and additional contributions to the manuscript. All authors read and approved the final submitted manuscript.

## Acknowledgements

We would like to thank the authors of Sharrar et al. [41] for performing sampling and sequencing, and for making the soil metagenomes and associated metadata publicly available. We acknowledge the use of MRC-CLIMB for the provision of high-performance servers, without which this work wouldn’t be possible.

---

**Fig. S1:**
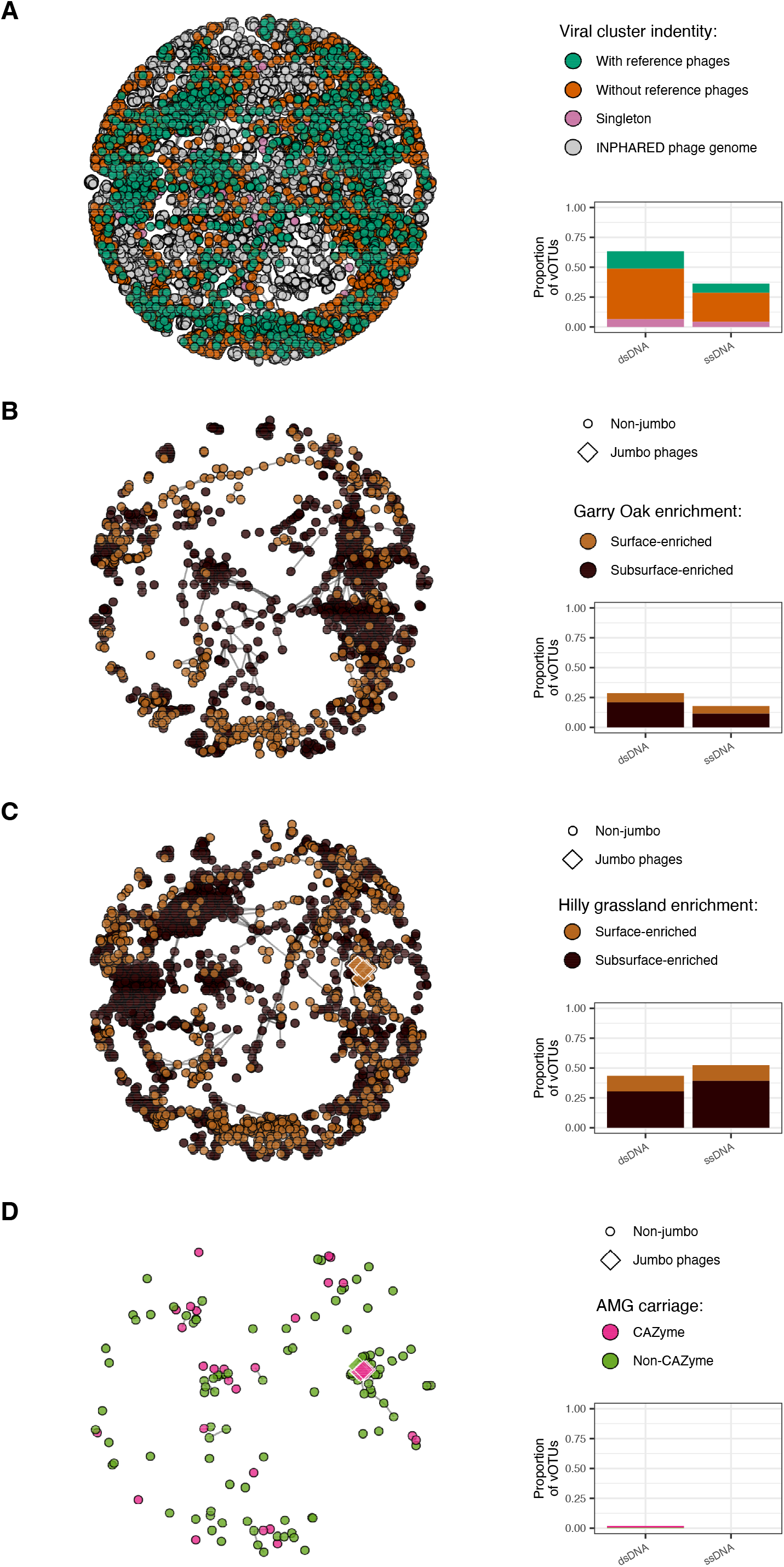
Taxonomic novelty of recovered soil vOTUs. Shared protein content of recovered soil vOTUs with previously discovered phage genomes. Network graph visualisations are annotated to represent **A** viral cluster identities (6124 dsDNA vOTUs, 193 ssRNA vOTUs, and 11,600 reference genomes), **B** depth enrichment in Garry Oak (1637 dsDNA vOTUs, 19 ssDNA vOTUs), **C** depth enrichment in Hilly grassland (2820 dsDNA vOTUs, 138 ssDNA vOTUs), and **D** vOTUs carrying AMGs (152 dsDNA vOTUs, 0 ssDNA vOTUs). Bar charts (right) summarise the proportion of dsDNA vOTUs and ssDNA vOTUs included in each network visualisation. Depth enrichment represents vOTUs enriched in either surface soil (20 cm) or subsurface soil (40 cm – 115 cm).

**Fig. S2:**
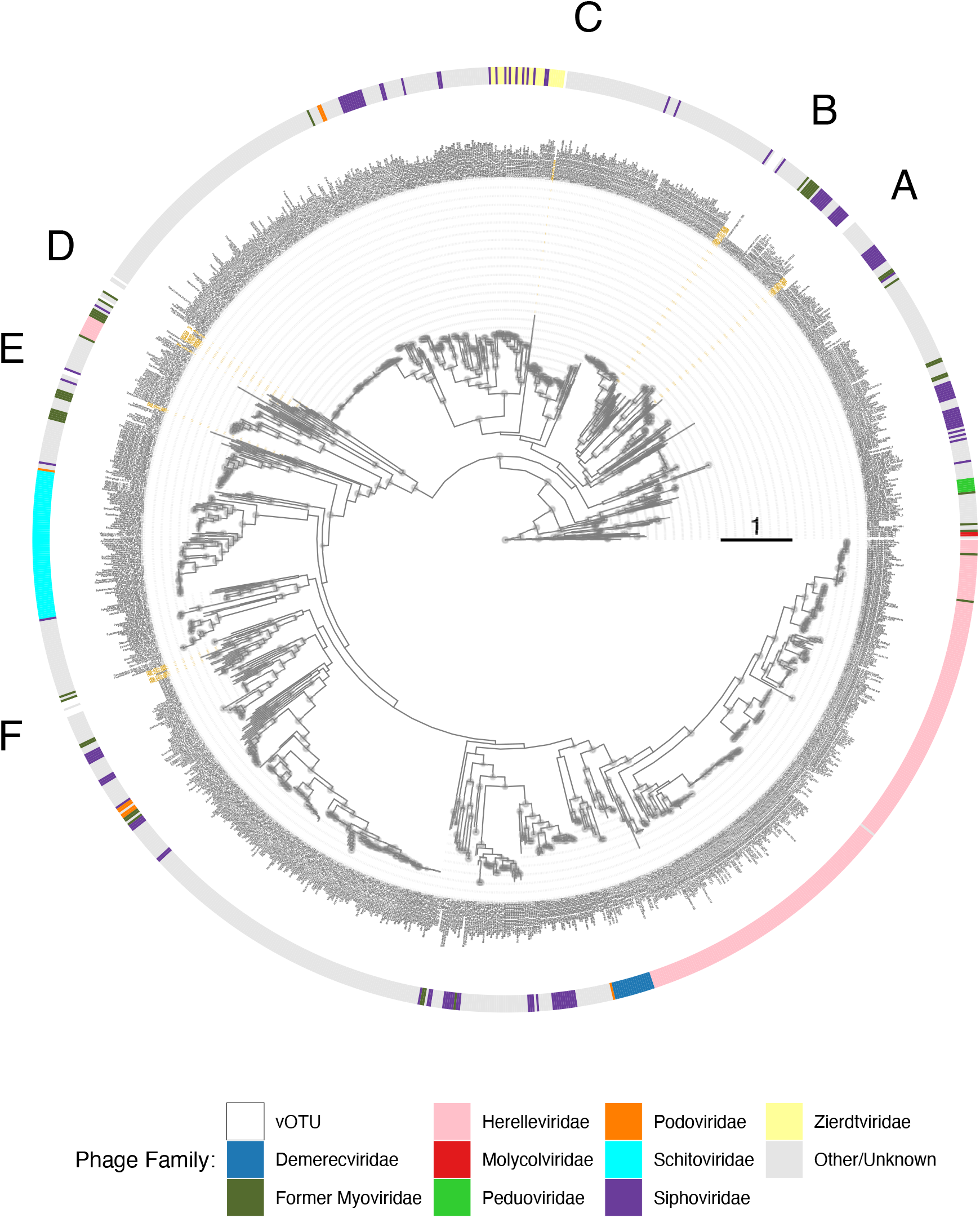
Phylogenetic assessment of jumbo phage vOTUs and jumbo-related vOTUs using DNA polymerase gene. Phylogeny of jumbo phage vOTUs and vOTUs sharing viral clusters with jumbo phage vOTUs (jumbo-related vOTUs) using translated DNA polymerase sequences. Phylogenetic tree contains 1284 DNA polymerase sequences from 1205 previously isolated phage sequences and 24 DNA polymerase sequences from 14 vOTUs recovered in this study (eight jumbo phage vOTUs and six jumbo-related vOTUs). Brand node labels indicate branch support: ≥ 0.9 (large circles), ≥ 0.8 (medium circles), ≥ 0.7 (small circles), < 0.7 (no circle). Tip labels indicate genome sequence name; vOTUs recovered in this study are labelled in gold. Outer ring fill colour denotes known phage families. Letters indicate the locations of 6 distinct phylogenetic groups of jumbo phage vOTUs and jumbo-related vOTUs.

**Fig. S3:**
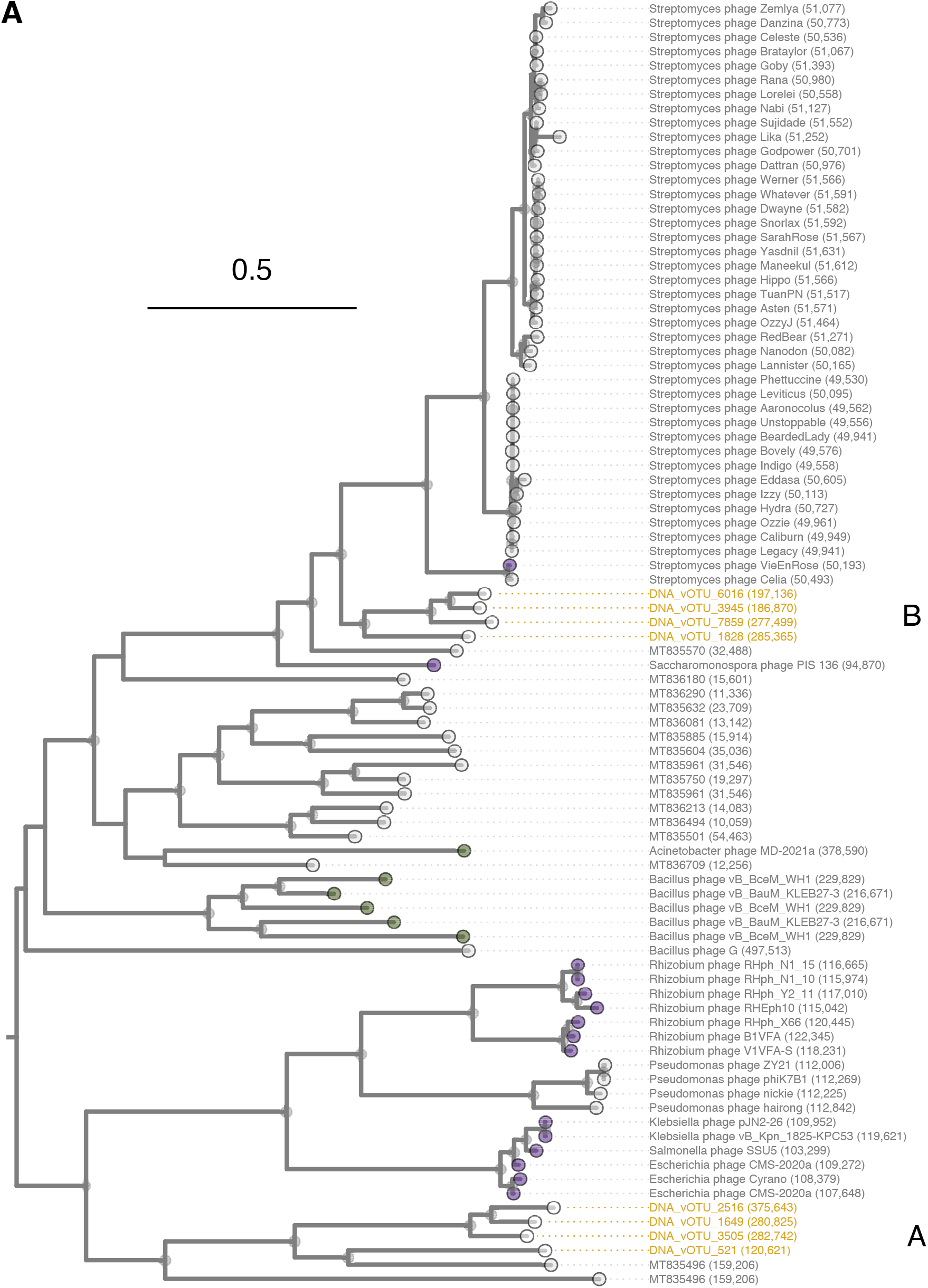
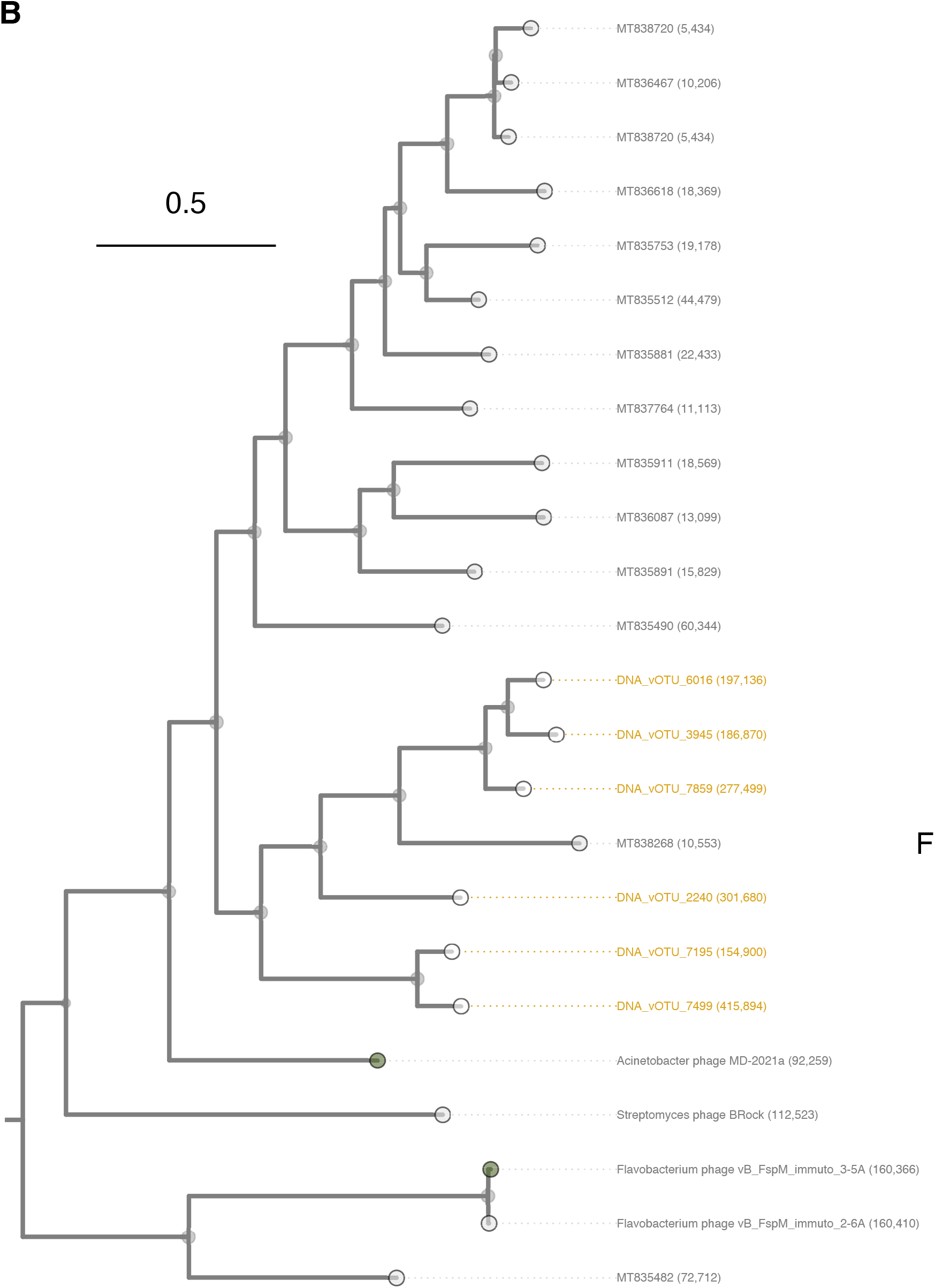
Phylogenetic groups A, B, and F from assessment of jumbo phage vOTUs and jumbo-related vOTUs using DNA polymerase gene. Further investigation of distinct phylogenetic groups identified from Fig. S2: **A** Groups A and B, **B** Group F. Brand node labels indicate branch support: ≥ 0.9 (large circles), ≥ 0.8 (medium circles), ≥ 0.7 (small circles), < 0.7 (no circle). Tip node fill colour denotes known phage families. Tip labels indicate genome sequence name and genome length in bp; vOTUs recovered in this study are labelled in gold. Letters indicate the locations of distinct phylogenetic groups of jumbo phage vOTUs and jumbo-related vOTUs.

**Fig. S4:**
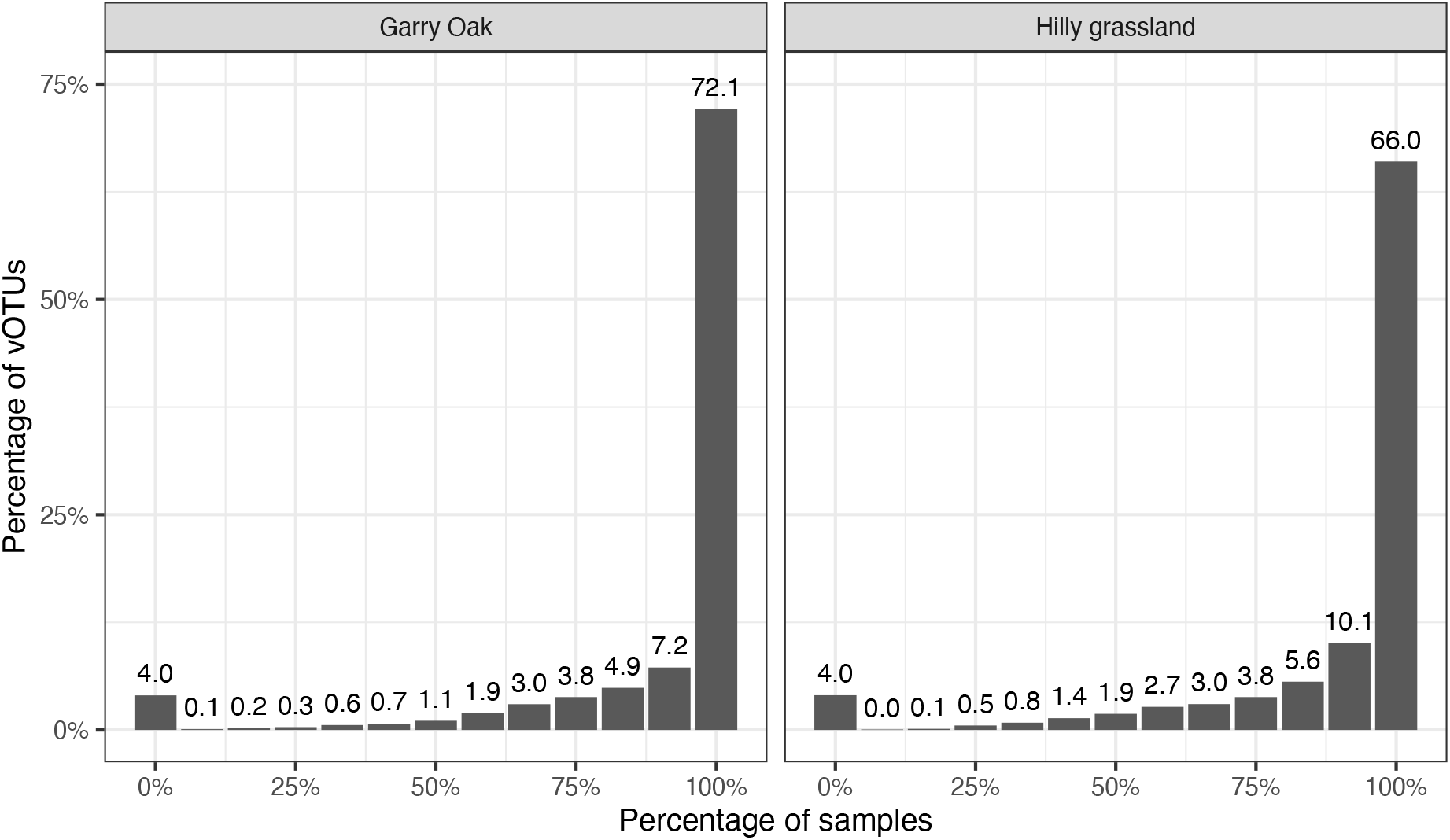
Prevalence of viral populations. Percentage of vOTUs detected in at each percentage of soil samples. Number above bars specify the percentage of vOTUs detected.

**Fig. S5:**
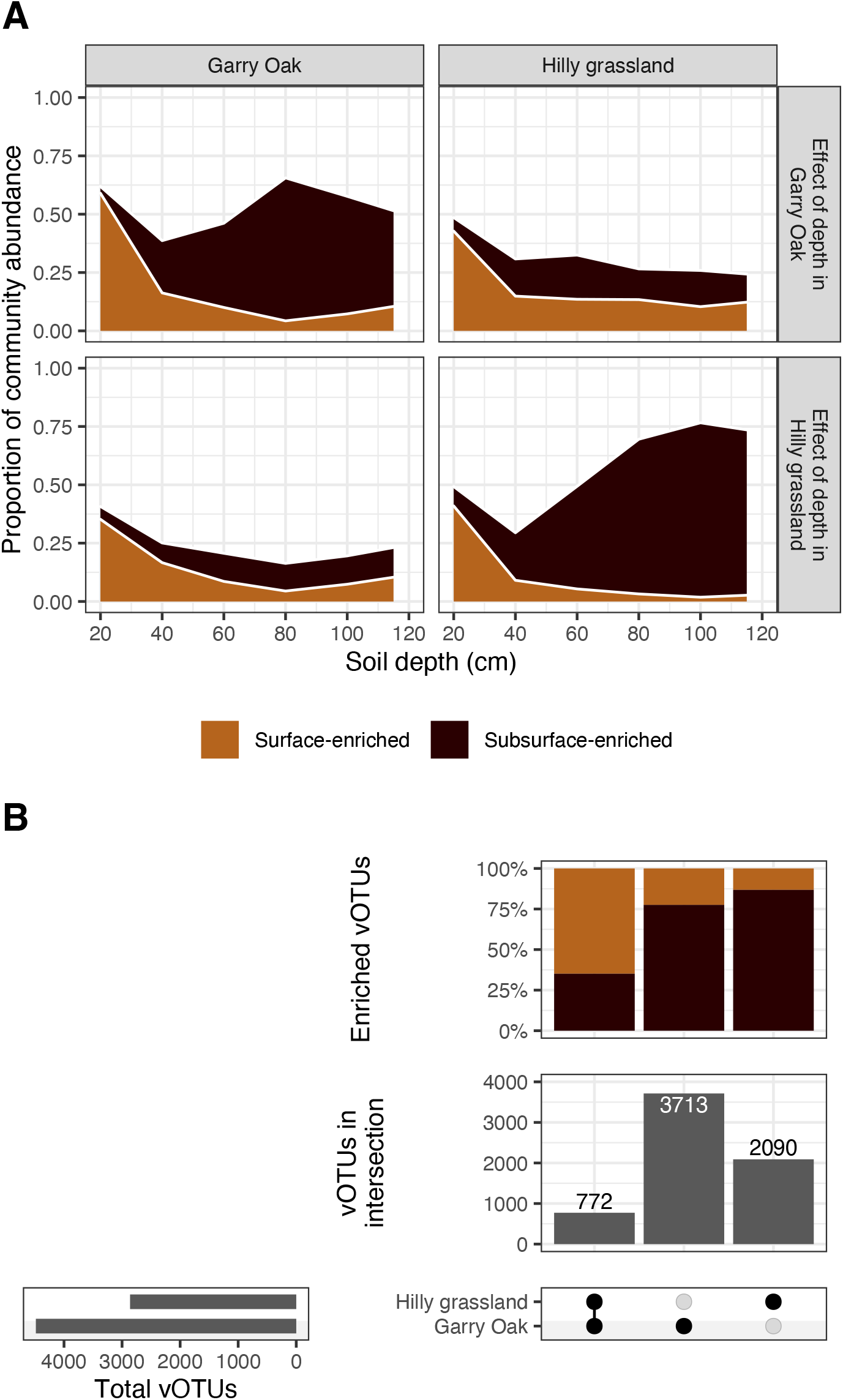
Overlap in depth-enrichment of viral populations between sites. **A** Relative abundance of depth-enriched viral populations. Proportional abundance of vOTUs enriched in either surface soil (20 cm) or subsurface soil (40 cm – 115 cm) based on samples derived from Garry Oak and Hilly grassland, across Garry Oak and Hilly grassland samples, respectively. Fill colour indicates enrichment: surface-enriched (light brown) or subsurface enriched (dark brown). **B** Overlap in depth enrichment of viral populations between sites. Intersection matrix denoting site investigated (bottom-right), total vOTUs detected in each site (bottom-left), number of enriched vOTUs in site intersection (middle-right), percentage of enriched vOTUs corresponding to surface-enriched or subsurface-enriched, respectively (top-right).

**Fig. S6:**
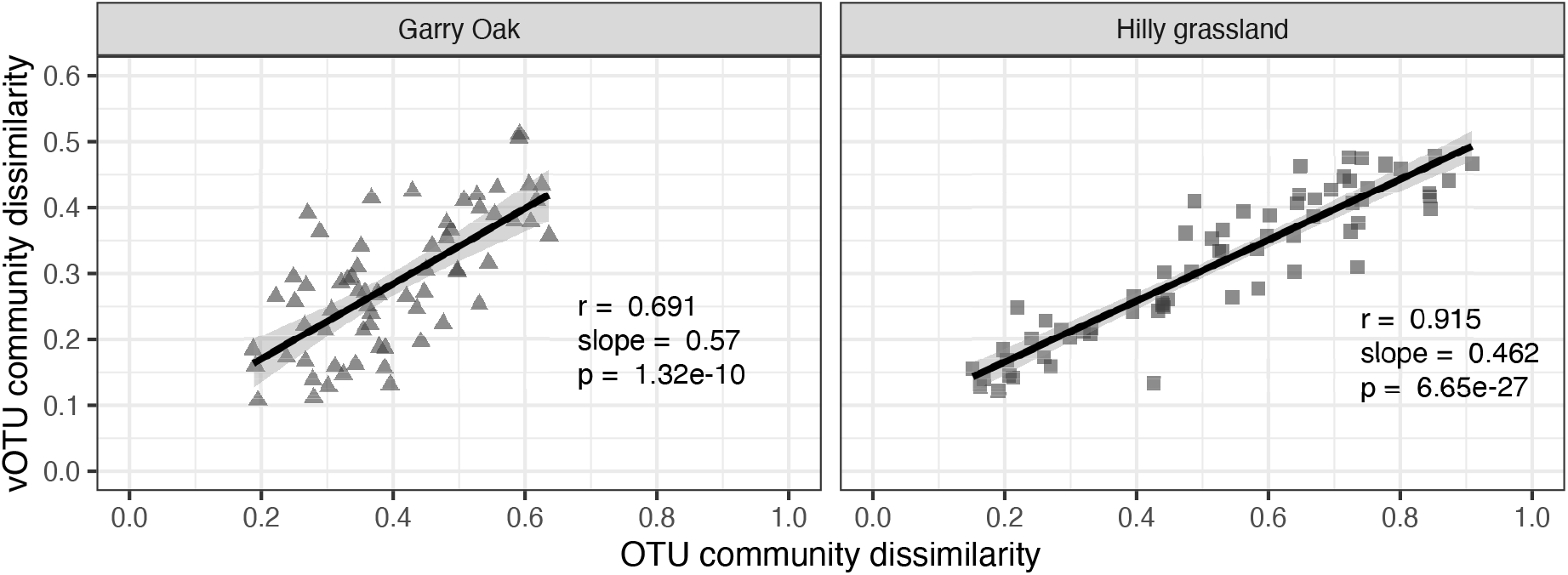
Correlation of viral community and microbial community structure. Trend lines represent linear regression estimates, with shaded cloud representing 95% confidence interval. *r* corresponds to Pearson’s correlation coefficient and *p* corresponds to the associated p-value.

**Fig. S7:**
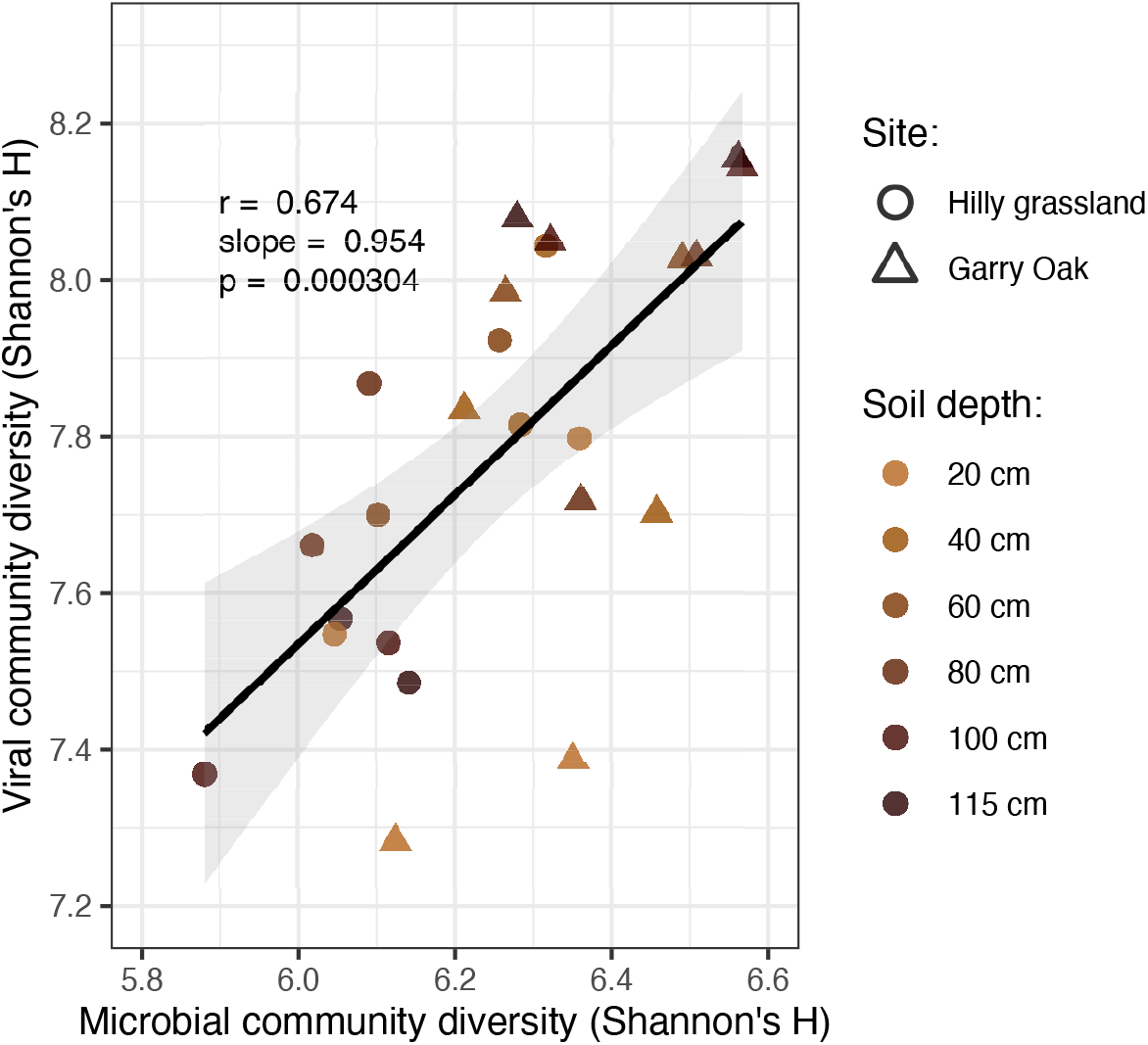
Correlation of viral community and microbial community diversity. Trend line represents linear regression estimates, with shaded cloud representing 95% confidence interval. *r* corresponds to Pearson’s correlation coefficient and *p* corresponds to the associated p-value. Shapes indicate site: Hilly grassland (squares) and Garry Oak (triangles). Shapes are coloured based on soil depth.

**Fig. S8:**
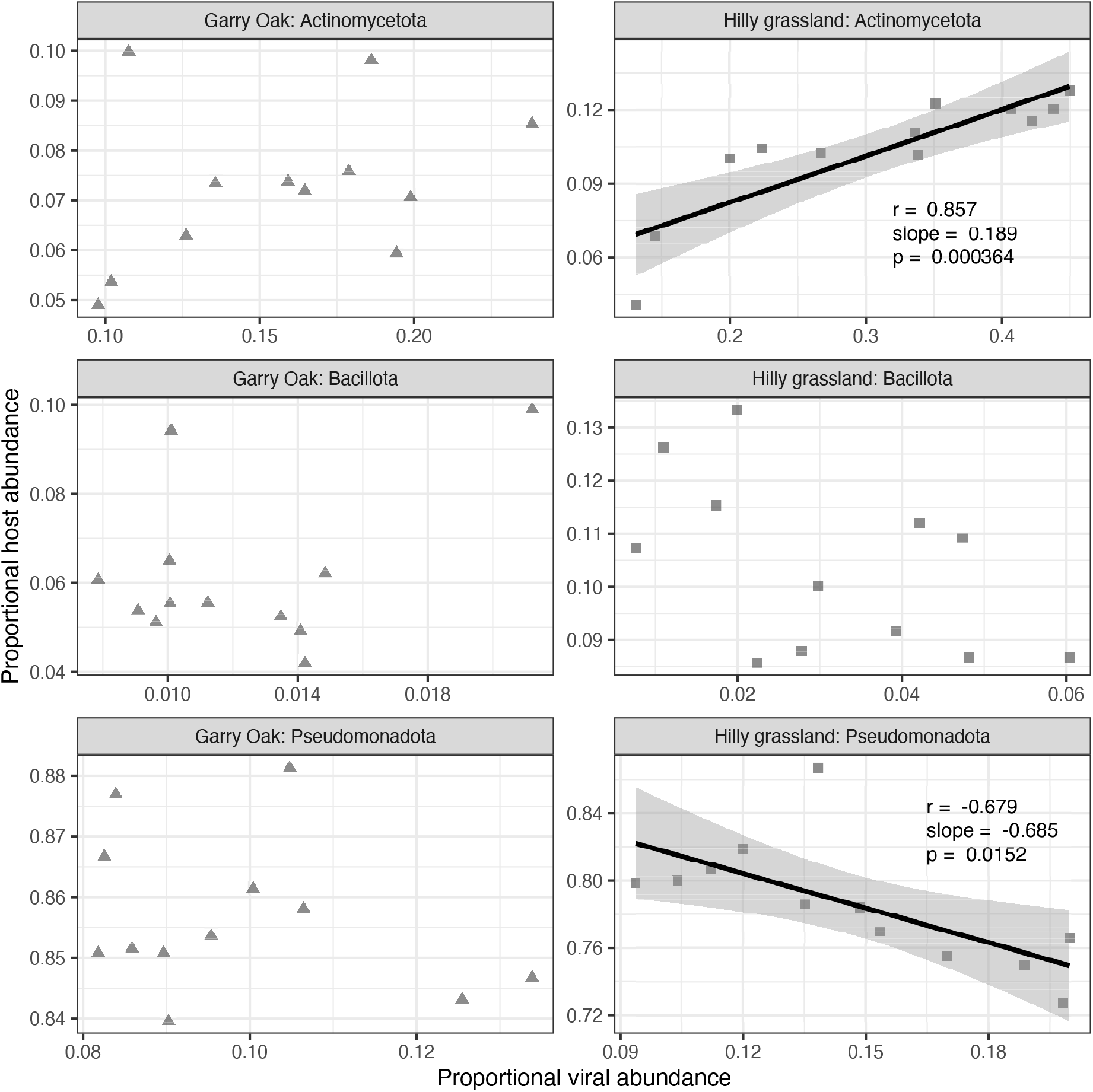
Correlation of viral abundances and host abundances. Trend line represents linear regression estimates, with shaded cloud representing 95% confidence interval. *r* corresponds to Pearson’s correlation coefficient and *p* corresponds to the associated p-value.

**Fig. S9:**
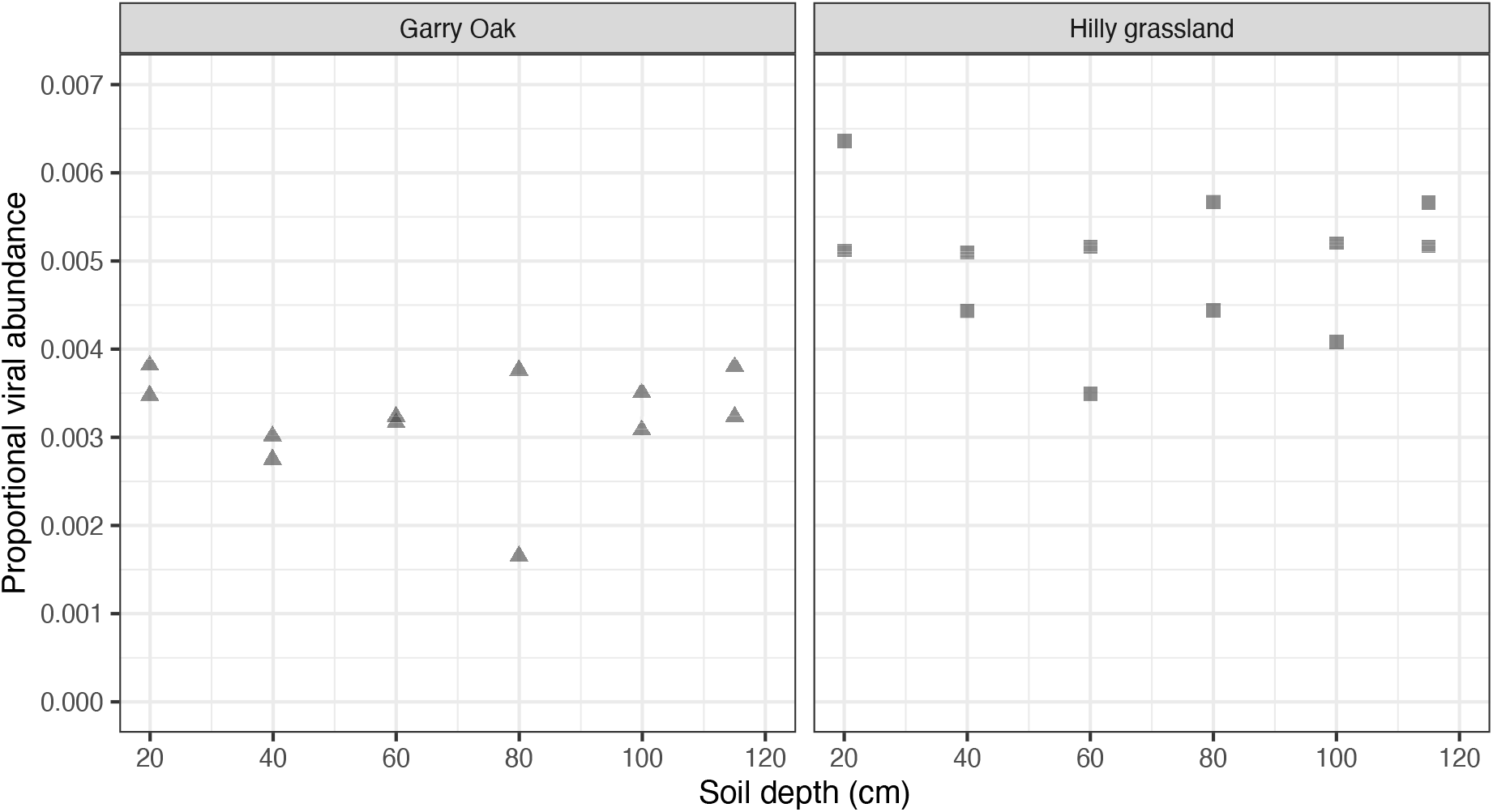
Relative abundance of viruses carrying carbohydrate-active enzymes.

**Fig. S10:**
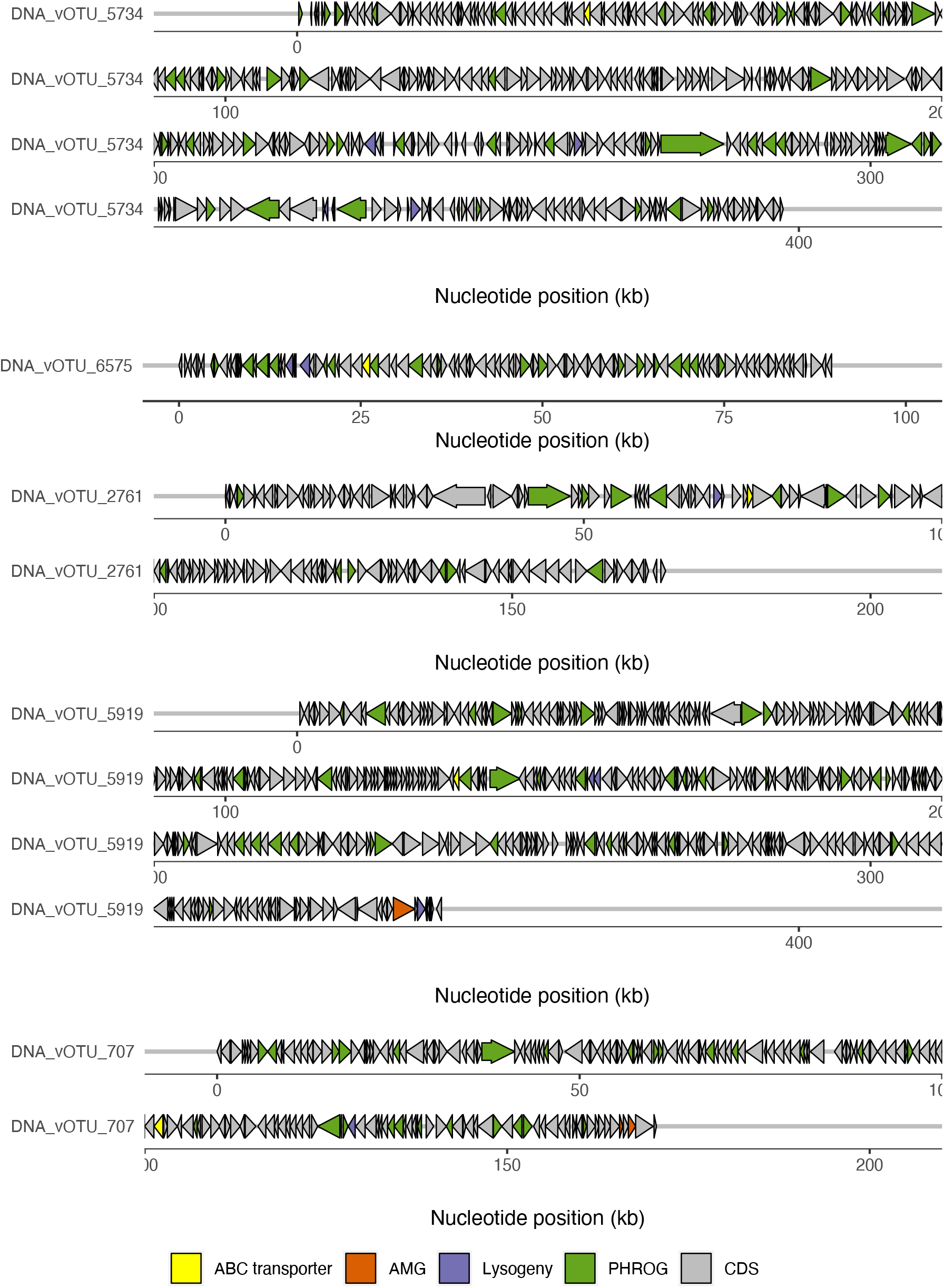
Genome maps of high-quality viral genomes carrying ABC transporters under positive selection. Arrow fill colour indicates gene function.

**Fig. S11:**
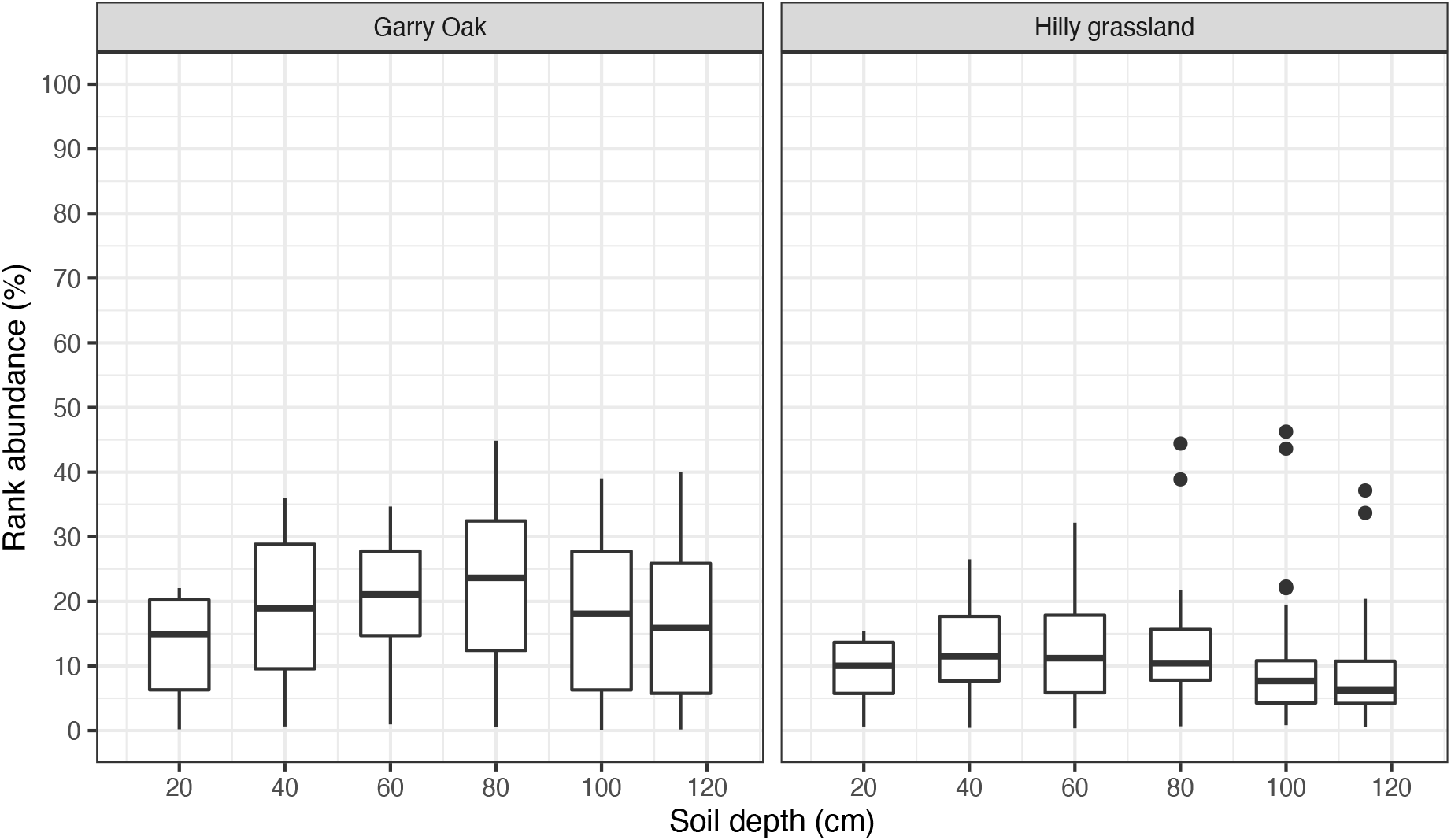
Rank abundance of jumbo phages. Rank abundance represented as a percentage of 10,196 vOTUs. 0% indicates the lowest rank and the most abundant vOTU, while 100% indicates the highest rank and the least abundant vOTU. Boxes denote median, upper, and lower quartiles. Whiskers indicate minimal and maximal values, with outliers in filled circles.

**Fig. S12:**
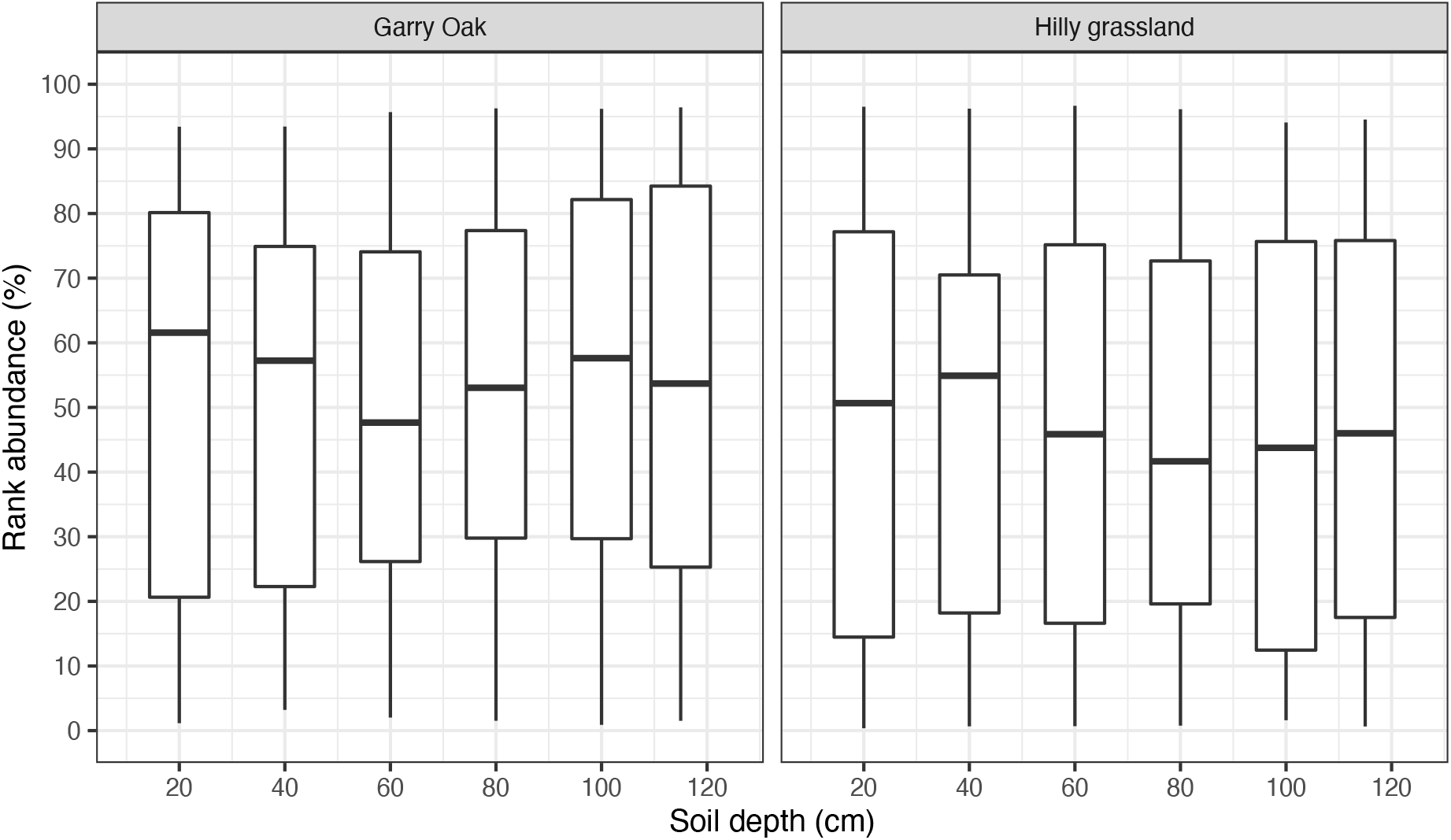
Rank abundance of CAZyme-carrying vOTUs. Rank abundance represented as a percentage of 10,196 vOTUs. 0% indicates the lowest rank and the most abundant vOTU, while 100% indicates the highest rank and the least abundant vOTU. Boxes denote median, upper, and lower quartiles. Whiskers indicate minimal and maximal values, with outliers in filled circles.

**Fig. S13:**
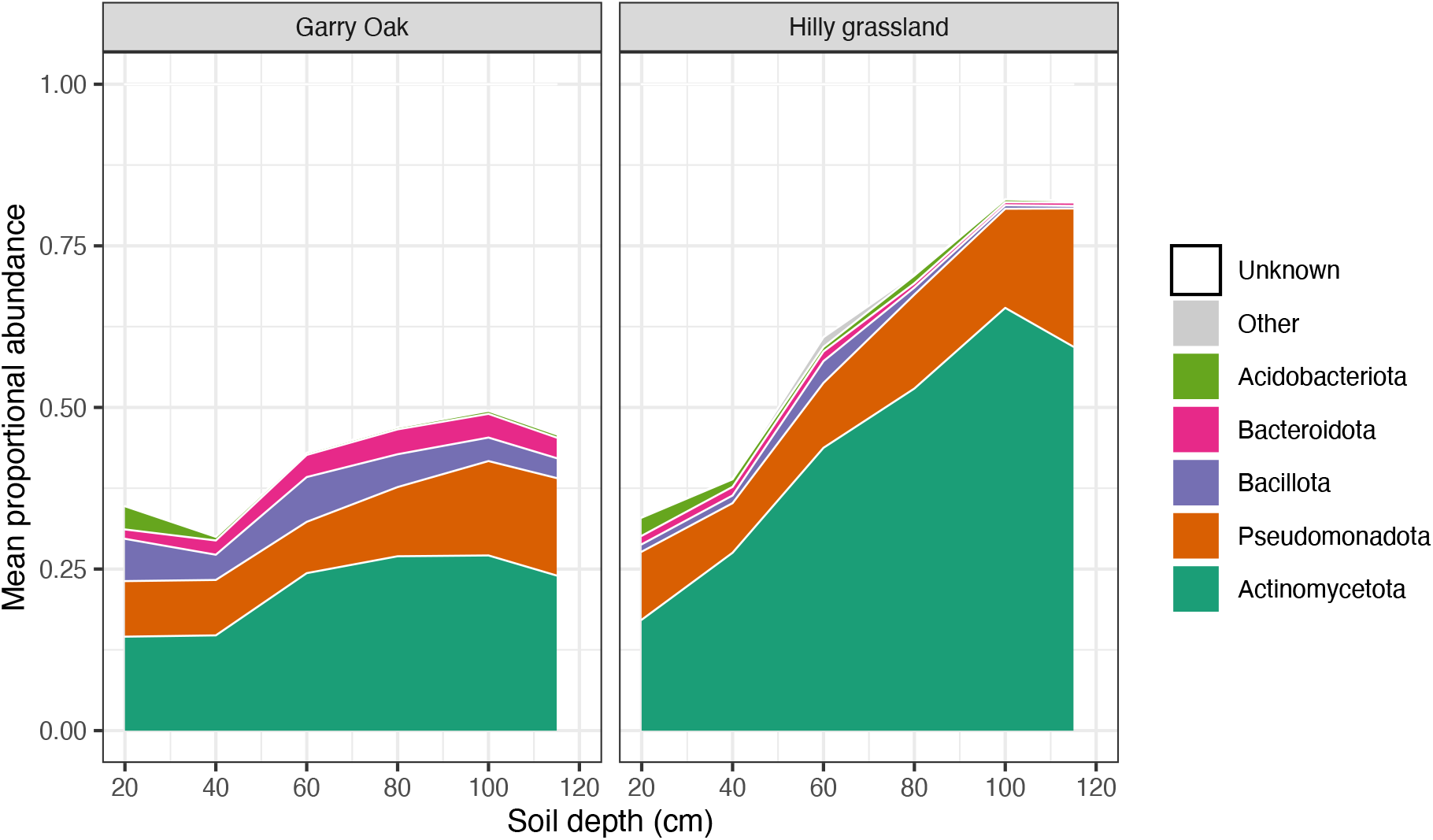
Relative abundance of hosts of viruses carrying auxiliary metabolic genes. Proportional abundance of vOTUs carrying AMGs plotted across soil depth. Fill colour indicates host phyla.

**Fig. S14:**
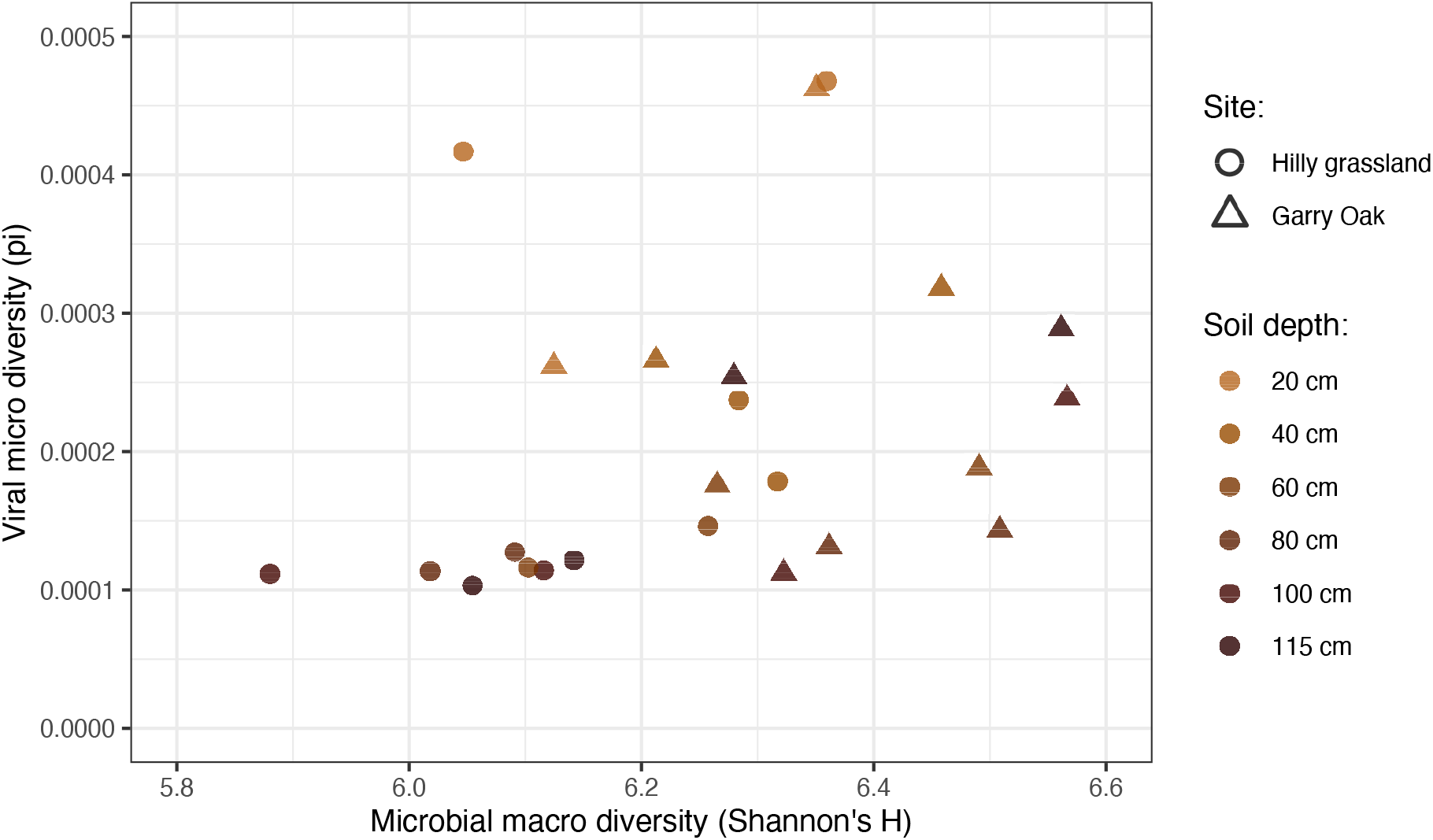
Correlation of microbial macro diversity and viral micro diversity. Shapes indicate site: Hilly grassland (squares) and Garry Oak (triangles). Shapes are coloured based on soil depth.

